# Steamed broccoli sprouts alleviate DSS-induced inflammation and retain gut microbial biogeography in mice

**DOI:** 10.1101/2023.01.27.522641

**Authors:** Johanna M. Holman, Louisa Colucci, Dorien Baudewyns, Joe Balkan, Timothy Hunt, Benjamin Hunt, Marissa Kinney, Lola Holcomb, Grace Chen, Peter L. Moses, Gary M. Mawe, Tao Zhang, Yanyan Li, Suzanne L. Ishaq

## Abstract

Inflammatory Bowel Diseases (IBD) are devastating conditions of the gastrointestinal tract with limited treatments, and dietary intervention may be effective, and affordable, for managing symptoms. Glucosinolate compounds are highly concentrated in broccoli sprouts, especially glucoraphanin, and can be metabolized by certain mammalian gut bacteria into anti inflammatory isothiocyanates, such as sulforaphane. Gut microbiota exhibit biogeographic patterns, but it is unknown if colitis alters these or whether the location of glucoraphanin metabolizing bacteria affects anti-inflammatory benefits. We fed specific pathogen free C57BL/6 mice either a control diet or a 10% steamed broccoli sprout diet, and gave a three-cycle regimen of 2.5% dextran sodium sulfate (DSS) in drinking water over a 34-day experiment to simulate chronic, relapsing ulcerative colitis. We monitored body weight, fecal characteristics, lipocalin, serum cytokines, and bacterial communities from the luminal and mucosa-associated populations in the jejunum, cecum, and colon. Mice fed the broccoli sprout diet with DSS treatment performed better than mice fed the control diet with DSS, including significantly more weight gain, lower Disease Activity Indexes, lower plasma lipocalin and proinflammatory cytokines, and higher bacterial richness in all gut locations. Bacterial communities were assorted by gut location, but were more homogenous across locations in the control diet + DSS mice. Importantly, our results showed that broccoli sprout feeding abrogated the effects of DSS on gut microbiota, as bacterial richness and biogeography were similar between mice receiving broccoli sprouts with and without DSS. Collectively, these results support the protective effect of steamed broccoli sprouts against dysbiosis and colitis induced by DSS.

**Importance:** Evaluating bacterial communities across different locations in the gut provides a greater insight than fecal samples alone, and provides an additional metric by which to evaluate beneficial host-microbe interactions. Here, we show that 10% steamed broccoli sprouts in the diet protects mice from the negative effects of dextran sodium sulfate induced colitis, that colitis erases biogeographical patterns of bacterial communities in the gut, and that the cecum is not likely to be a significant contributor to colonic bacteria of interest in the DSS mouse model of ulcerative colitis. Mice fed the broccoli sprout diet during colitis performed better than mice fed the control diet while receiving DSS. The identification of accessible dietary components and concentrations that help maintain and correct the gut microbiome may provide universal and equitable approaches to IBD prevention and recovery, and broccoli sprouts represent a promising strategy.

## Introduction

Inflammatory Bowel Diseases (IBD) are globally prevalent, chronic inflammatory diseases of multifactorial origin which disrupts daily life and creates financial burdens to individuals and healthcare systems (1, 2). IBD symptoms occur in the gastrointestinal (GI) tract, and can be accompanied by immune dysfunction (3) and microbial community changes in the gut. In addition to being debilitating, a longer duration of IBD is associated with an increased risk of developing GI cancers (4). Treatments are currently limited to alleviating inflammation and returning patients to as close to homeostasis as possible. Diet can play an important role in the management of IBD as a source of anti-inflammatory metabolites, and as a tool for influencing the robustness of gut microbiomes. However, many guidelines for IBD patients recommend avoiding high-fiber or sulfur-rich foods which produce metabolites that could exacerbate symptoms (5). Diet can be beneficial (6) or detrimental to gut inflammation (7), but IBD patients may choose to avoid most fiber or sulfur-rich foods rather than test their reaction to specific foods (8). Thus, a better understanding of the interaction of diet, gut microbiota, and disease is needed before dietary recommendations can be made.

Diets which are high in cruciferous vegetables, such as broccoli, are associated with reduced inflammation and cancer risk, e.g. (9–11), due to a group of specific plant secondary compounds, glucosinolates. Glucosinolates are very high in broccoli, especially seeds or immature sprouts, and can be converted into bioactive metabolites (12, 13), such as sulfur-containing isothiocyanates, by the action of myrosinase, an enzyme present in these vegetables. Isothiocyanates are used by the plant for defense against insect herbivory, but in humans, have been identified as bioactive candidates for reducing gut inflammation (14–16). Specifically, sulforaphane (SFN), a well-studied isothiocyanate (17), has been shown to inhibit the action of immune factors which are responsible for upregulation of several proinflammatory cytokines; interleukins-6, -8, -12, -21, and -23 (11, 18, 19), which has recently been evaluated as a possible strategy for reducing gut inflammation in humans and mouse models (13, 14, 16).

However, consuming raw broccoli and broccoli sprouts results in small amounts of available SFN, as broccoli-sourced enzymes preferentially metabolize the precursor, glucoraphanin (GLR), to an inactive byproduct instead (17). Cooking the sprouts alters the activity of plant enzymes to prevent the creation of the inactive byproduct, leaving glucosinolates intact (13, 20). Mammals do not produce the enzymes for converting glucosinolates to isothiocyanates; however, gut bacteria with β-thioglucosidase activity can cleave the glycoside moiety from glucosinolates, and there is evidence for GLR hydrolysis to SFN by colonic and cecal bacteria, *ex vivo* and *in vivo* in humans and animal models, e.g. (13, 15, 17). The bacteria and genes that are capable of metabolizing glucosinolates are not fully known, but strains of *Lactobacillus*, *Bifidobacteria, Bacteroides,* and other genera have been implicated (15, 21, 22), with the most known about the genes in *Bacteroides thetaiotaomicron* (*21*). Moreover, the conditions in the gut (e.g., diet, inflammation) under which this glucosinolate metabolism by bacteria occurs or not are not well understood. For example, people consuming cooked broccoli stopped excreting isothiocyanates in their urine after being treated with oral antibiotics and bowel cleansing which reduced gut bacterial diversity and biomass (23). Further, not everyone’s gut bacteria can be reliably induced to meaningful levels of GLR conversion (24, 25). Thus, bacterial metabolism of GLR to SFN *in situ* must be better understood prior to making dietary recommendations.

Mouse studies suggest that biotransformation of GLR to SFN occurs in the colon (13, 21, 26), and we confirmed that the majority of SFN accumulation occurs in the colon by using multiple locations of measurement along the GI tract (13). The microbial communities along the GI tract are highly dependent on the diet, health status, age, and microbial encounters of the host (27, 28). Additionally, each organ in the GI tract and sites within organs foster different environmental conditions that create spatial niches for different microbial taxa, an ecological pattern known as biogeography (29, 30). However, changes to conditions in one location may have repercussions downstream. For example, a low pH in the stomach results in low bacterial diversity and biomass in the duodenum (31). IBD patients often have a less acidic GI tract (32) due to altered diet or treatments, and this allows more bacteria to survive transit through the stomach which may result in small intestinal bacterial overgrowth (33), especially in mucosal-associated fractions (34–36). Moreover, the bacterial communities in the colon of IBD patients are disrupted (37), but this may result from inflammation and disruption to host-microbial relations in the colon itself or from further upstream (34, 38). Thus, changing the biogeography of gut bacterial communities in IBD patients could alter the dynamics of SFN production, which could improve or lessen the benefits to the host of having this anti-inflammatory produced in the gut.

Steamed broccoli sprouts provided in the diet of mice has been demonstrated to reduce inflammation in mice with chemically induced colitis (13); however, there are significant knowledge gaps regarding the effect of broccoli sprouts on the gut microbiota. In particular, there are gaps regarding bacterial biogeography in general and that of the glucosinolate-metabolizing taxa specifically, and how this may impact health benefits to the host. To address these, we assessed the impact of steamed broccoli sprouts on the biogeographic pattern of gut microbiota and disease outcome in a mouse model of chronic, relapsing colitis. We fed specific pathogen free (SPF) C57BL/6 mice either a control diet or a 10% steamed broccoli sprout diet balanced by macronutrient and by fiber, starting at 7 weeks of age and continuing for another 34 days. The fresh broccoli sprouts were steamed to inactivate plant enzymes, and thus the metabolism of glucosinolates to isothiocyanates would rely on the gut microbiota. A three-cycle regimen of dextran sodium sulfate (DSS) in drinking water was given to stimulate chronic colitis in the mice, which is a well-established method for modeling ulcerative colitis (39–41). We analyzed the bacterial communities from the luminal-associated (i.e., digesta contents) and mucosal-associated (i.e., epithelial scrapings) fractions in the jejunum, cecum, and colon using 16S rRNA gene sequencing, and correlated these communities with metrics of disease activity including weight gain/stagnation, fecal characteristics, lipocalin (as a surrogate marker for inflammation in IBD), and proinflammatory cytokines (Figure 1).

**Figure 1.**
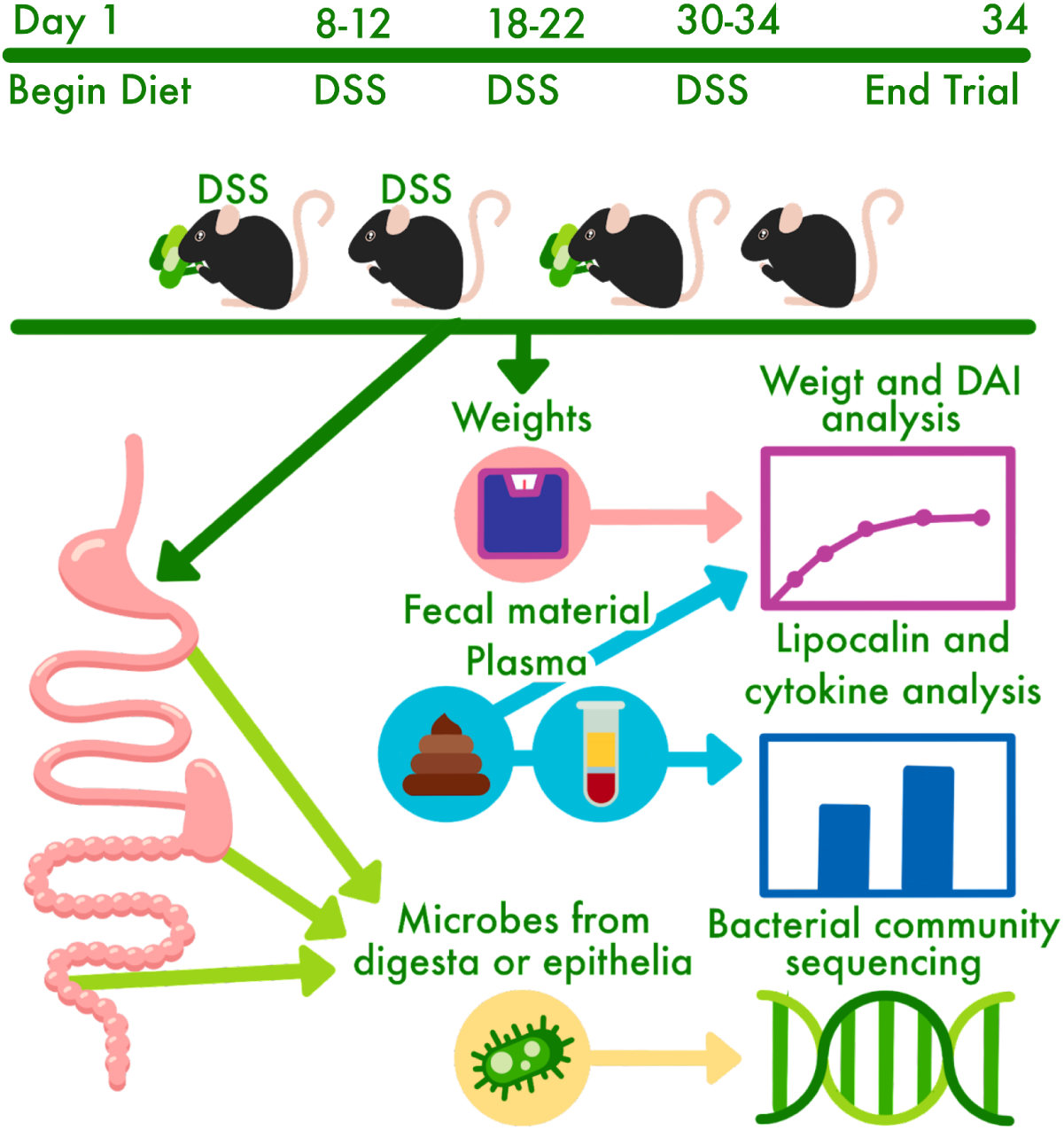
Experimental design schematic for a chronic model of ulcerative colitis induced by dextran sodium sulfate (DSS) in 40 male mice (C57BL/6) beginning at 7-weeks of age.

## Results

### Broccoli sprouts alleviated disease characteristics of DSS-induced colitis

After 1 week of acclimatization to the facility, forty (40) 7-week-old SPF C57BL/6 mice were divided into four groups: control diet (Control), control diet with 2.5% DSS in drinking water (Control+DSS), 10% (w/w) steamed broccoli sprout diet (Broccoli, with the control diet as the base), and 10% broccoli sprout diet with 2.5% DSS in drinking water (Broccoli+DSS). Over 34 days of the diet treatments, the two groups of mice receiving DSS were exposed to three cycles (days 8-12, 18-22, 30-34) designed to induce chronic, relapsing colitis (48, Figure 1). Periodically throughout the experiment, mice were monitored for weight loss, fecal blood, and fecal consistency. After euthanasia (day 34), samples were collected for analyses including: plasma lipocalin, plasma cytokines, and DNA extraction and community sequencing of lumen-associated (digesta contents) and mucosal-associated (epithelial scrapings) from the jejunum, cecum (contents only), and colon.

#### Weight

Mice with induced colitis and fed the broccoli sprout diet (Broccoli+DSS) gained significantly more weight than the mice with induced colitis on the control diet (Control+DSS) (Figure 2A; ANOVA, p < 0.03). Further, the weight gain of Broccoli+DSS mice was statistically similar to that for the two diet groups without DSS treatment (Figure 2A; ANOVA, p > 0.05). All the juvenile mice gained some weight as they were in a growth phase; despite this, the first two DSS cycles resulted in weight loss, especially in the Control+DSS group. The Broccoli+DSS mice recovered this weight loss prior to the beginning of the next cycle, but the Control+DSS mice did not (Figure 2A). At the beginning of the third and last DSS cycle, the mice were reaching maturity at eleven (11) weeks old and the weights of all groups were somewhat constant. However, the Control+DSS group demonstrated a significantly lower weight at the end of the study than the other groups which were statistically similar (Figure 2A).

**Figure 2.**
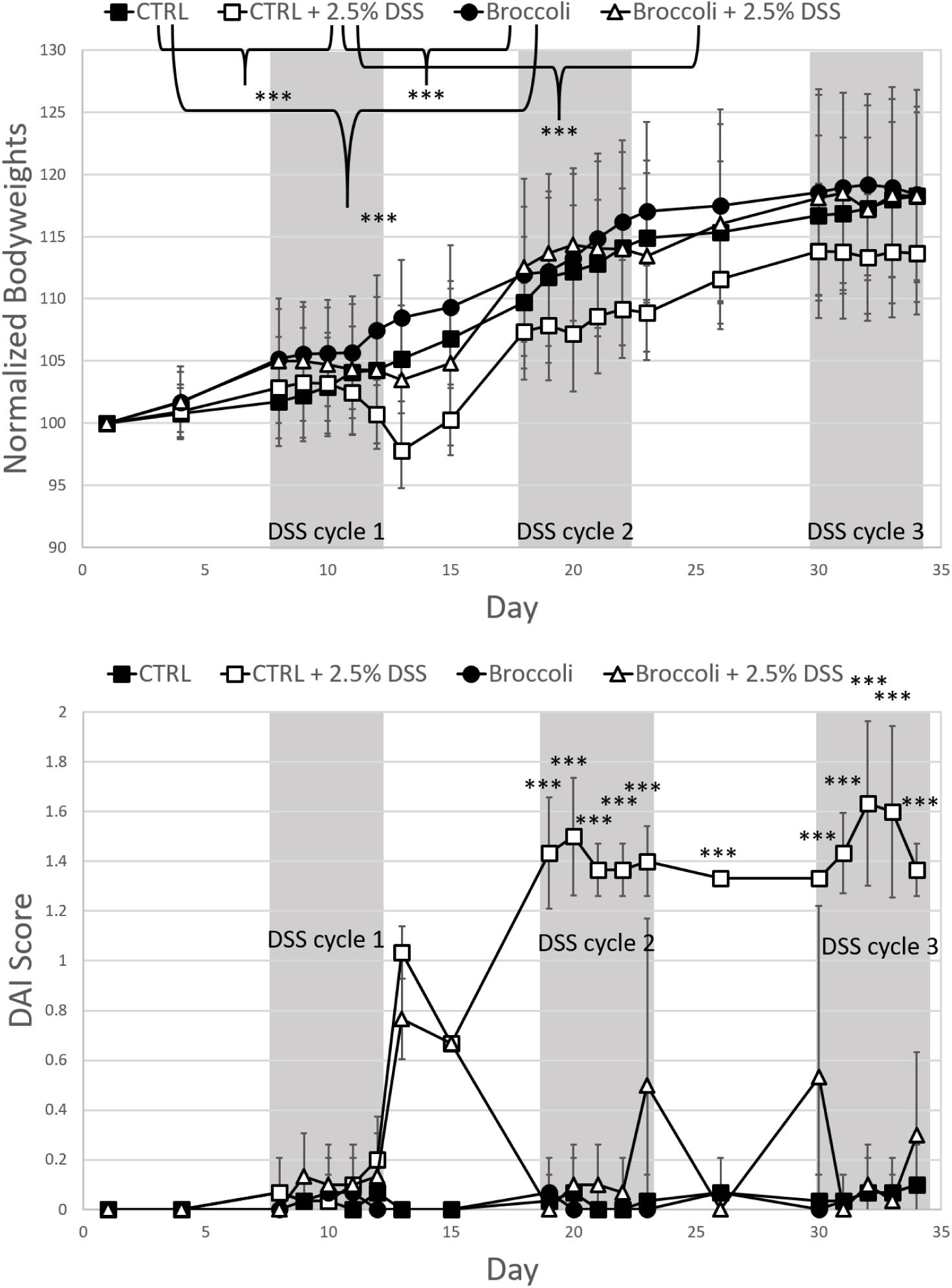
Body weights (A) and Disease Activity Index scores (B) in mice across a 34-day trial under a DSS-induced model of chronic, relapsing colitis with or without 10% steamed broccoli sprouts in diet. Body weights and time scale were normalized to the day mice began the broccoli sprout diet, set at 100% starting weight and Day 1, respectively. DAI scores are calculated by weight loss intensity score, fecal blood, and fecal consistency. Treatment comparisons at each day compared by ANOVA, and significance is designated as 0.001 ‘***’.

#### Disease Activity Index

Mice in the Broccoli+DSS group demonstrated significantly lower Disease Activity Index scores (DAI; including scoring weight loss, fecal blood, and fecal consistency) than the mice in the Control+DSS group during DSS cycle 2 and 3, but were similar in cycle 1 (Figure 2B). The Broccoli+DSS DAI scores were significantly elevated on four days during the trial (see Figure 2B; ANOVA, p < 0.05, comparisons by treatment, adjusted with TukeyHSD for multiple comparisons), as well as in an additive model considering the data over the entire study (GAM, F = 17.47, p < 0.001), however returned to near 0 during rest periods between DSS cycles. Mice in the Control+DSS group demonstrated elevated DAI scores from DSS cycle 1 and remained high throughout the study (Figure 2B), including during rest periods. The elevated scores were significant on many dates (see ***, Figure 2B) as compared to the mice in the Control group (Figure 2B, ANOVA, p < 0.05, comparisons by day, adjusted with TukeyHSD for multiple comparisons), as well as in an additive model considering the data over the entire study (GAM, F = 308.32, p < 0.001). The Control and Broccoli diet groups had statistically similar scores which were at or nearly 0, at the end of the trial (ANOVA, p > 0.05) or over time (GAM, p > 0.05).

#### Proinflammatory cytokines

Mice in the Broccoli+DSS group indicated significantly lower levels of proinflammatory cytokines IL-1β, IL-6 and TNF-α in plasma than mice in the Control+DSS group (Figure 3A). The plasma levels of IL-1β and TNF-α were similar across the Broccoli+DSS, Broccoli, and Control groups, and the IL-6 measure was slightly elevated (Figure 3A). The cytokine CCL4, also known as macrophage inflammatory protein 1β (MIP1β) and which is a chemokine that plays a critical role in regulating the migration and activation of immune cells during inflammatory processes (42), did not differ significantly among the four treatment groups (Figure 3A).

**Figure 3.**
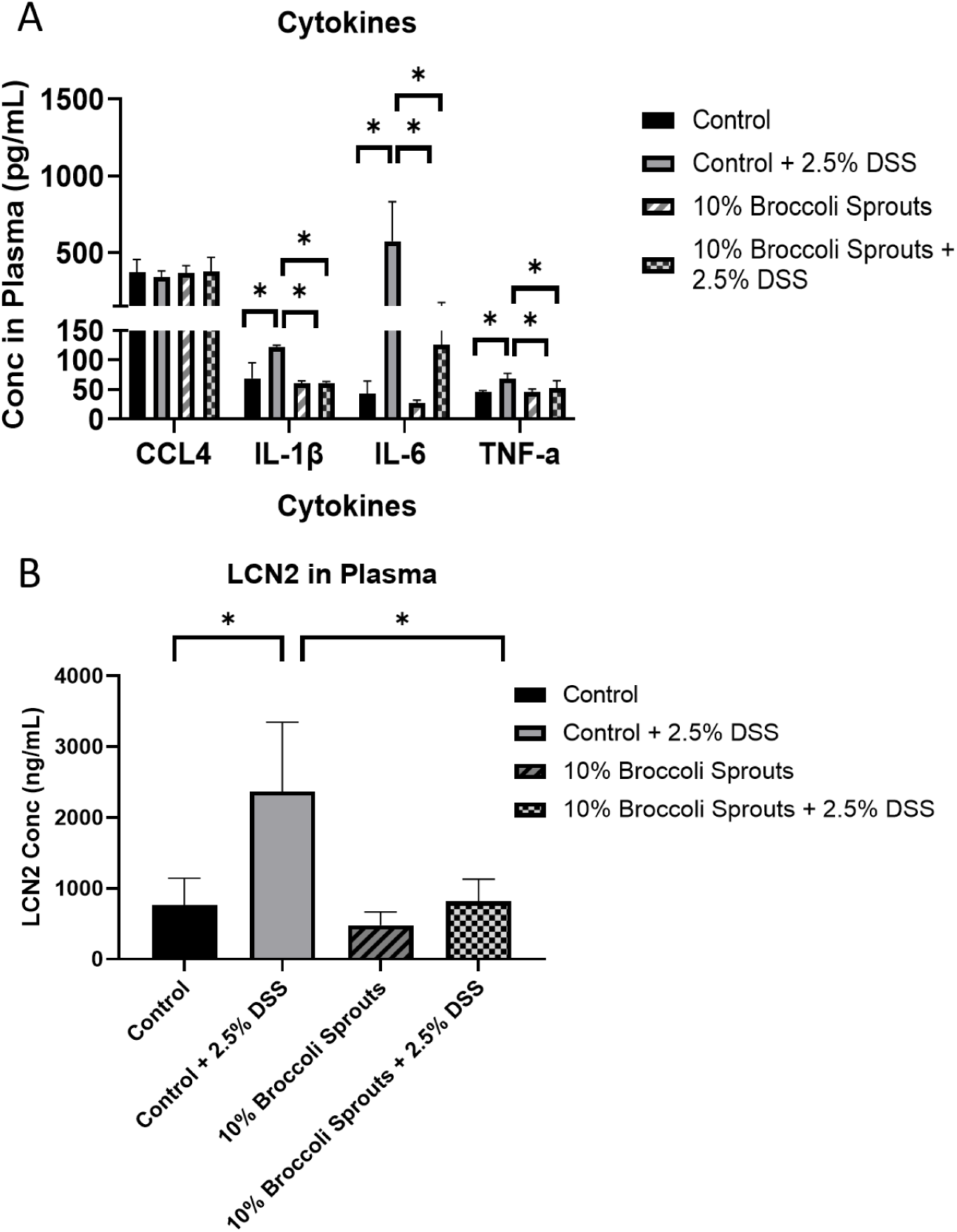
Cytokines (A) and lipocalin (B) in plasma on the last day of the trial from mice under a DSS-induced model of chronic, relapsing colitis with or without 10% steamed broccoli sprouts in diet. Cytokines include CCL4; interleukins IL-1β and IL-6; and tumor necrosis factor alpha (TNF-α). Data are represented as mean ᑊ± SD. *, p < 0.05, ANOVA.

#### Plasma lipocalin

At the end of DSS cycle 3, plasma lipocalin (LCN2) levels in the Broccoli+DSS mice were not significantly different from the groups without DSS, and were significantly lower in comparison with the Control+DSS group (ANOVA, p < 0.05, Figure 3B).

#### Fecal blood correlated with bacterial taxa

Mice in the Broccoli+DSS group with positive fecal occult blood scores demonstrated some differential bacteria compared to mice in the Control+DSS group with positive blood scores (Figure 4). The most abundant of these differential bacterial sequence variants (SV) in the Broccoli+DSS group were in the Muribaculaceae family and the *Clostridium* sensu stricto [Latin: strictly speaking] clade (Figure 4). The Control+DSS mice indicated different SVs with the most abundant in the *Clostridium* sensu stricto clade and *Cutibacterium* (Figure 4).

**Figure 4.**
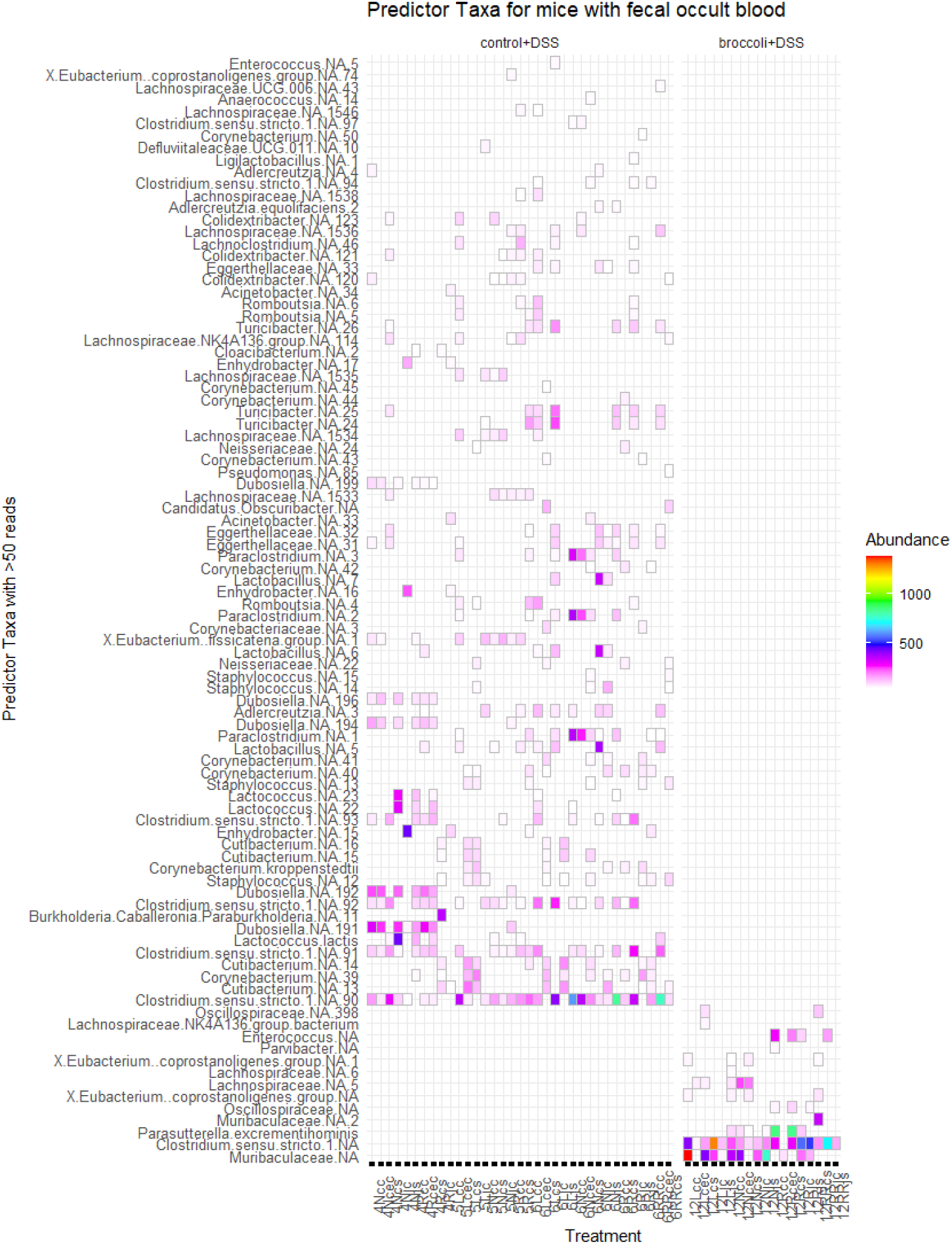
Gut bacterial taxa identified from mice having a positive fecal occult blood score at sacrifice under a DSS-induced model of chronic, relapsing colitis with or without 10% steamed broccoli sprouts in diet. Data were subset to mice with a positive score on the last day of the experiment, when the gut samples were collected. Important features (SVs) were identified through permutational random forest analysis, and only the features important to this group (>50 reads) are listed out of 143 significant (p < 0.05) features across all treatments. Model accuracy was 89.7%. Bacterial sequence variants (SV) are identified as the lowest level of taxonomic identity possible, with “NA” indicating which could not be identified to species, and the number indicating which specific SV it was.

### Broccoli sprouts protected against DSS-induced changes to bacterial communities

#### Bacterial richness

Mice with induced colitis on the broccoli sprouts diet (Broccoli+DSS) demonstrated significantly more bacterial richness in the cecal contents, colon contents, and colon scrapings than mice without broccoli sprouts (Control+DSS, Figure 5, Table 1). Across all treatments and all anatomical sites (jejunum contents & scrapings, cecal contents, colon contents & scrapings), bacteria were primarily identified as belonging to the phyla Firmicutes/Bacillota, Bacteroidetes/Bacteroidota, Proteobacteria/Pseudomonadota, Actinobacteria/Actinomycetota (which was highest in Control+DSS mice), and Verrucomicrobia/Verrucomicrobiota (which was highest in mice consuming broccoli sprouts; (Figure S1). The two groups receiving broccoli sprouts had the highest bacterial richness in all gut locations, particularly in the colon contents and scrapings (Figure 5). In comparison to the Control group, the Control+DSS mice demonstrated higher bacterial richness in the jejunum, lower richness in the cecal contents, and comparable bacterial richness in the colon (Figure 5). In addition, the Control+DSS mice had similar median richness in the jejunum, cecum, and colon contents, with the widest distribution of richness in the jejunum and colon (Figure 5). Moreover, the Control+DSS richness was similar between the digesta contents and epithelial scraping sites, in both the jejunum and in the colon, implying homogenization of bacterial communities in those sites (Figure 5).

**Figure 5.**
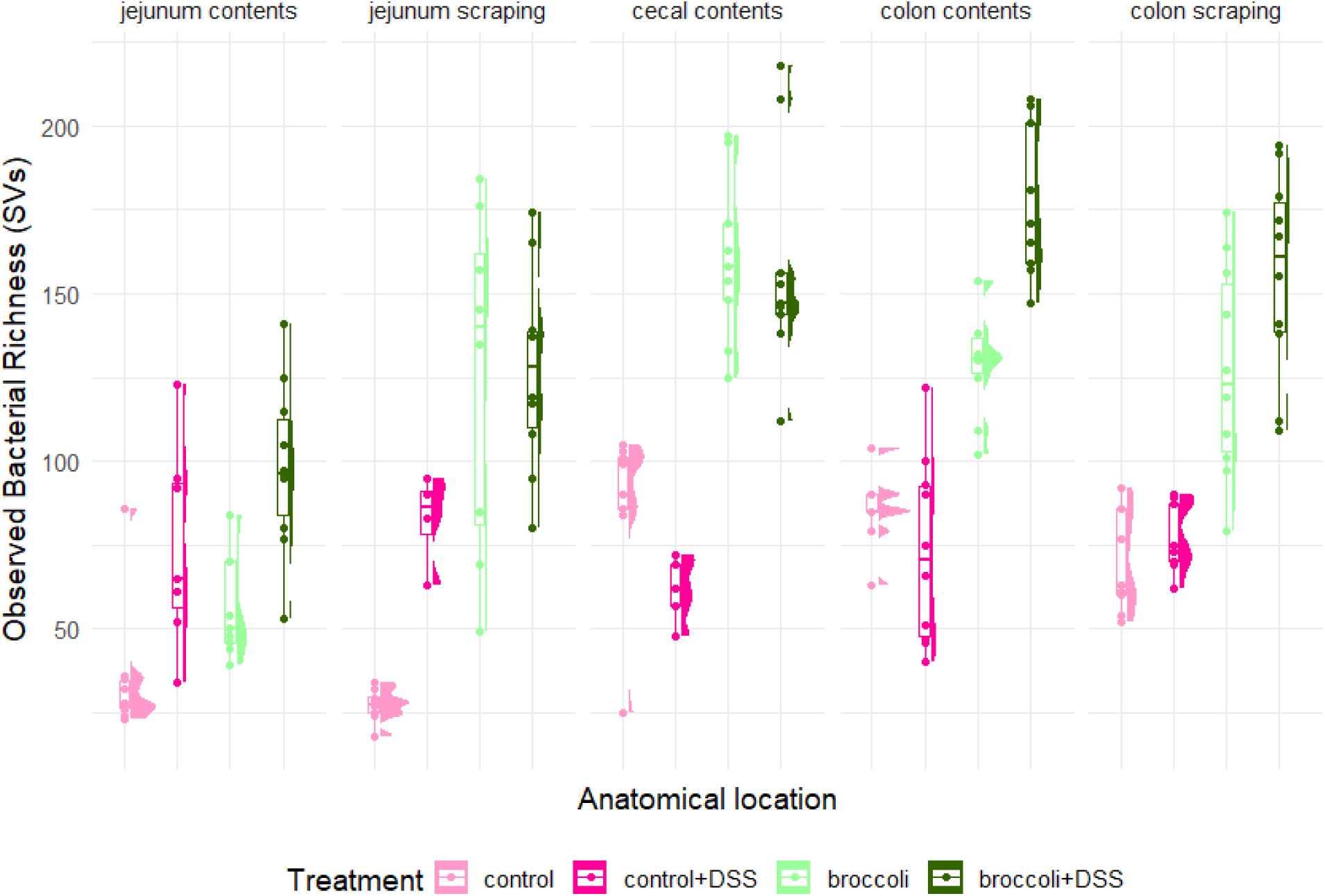
Observed bacterial richness along the intestine of mice under a DSS-induced model of chronic, relapsing colitis with or without 10% steamed broccoli sprouts in diet. Richness is calculated as the number of different bacterial sequence variants (SVs). Statistically significant comparisons are provided in Table 1.

**Table 1.** Statistical comparison of observed richness along the intestine of mice under a DSS induced model of chronic, relapsing colitis with or without 10% steamed broccoli sprouts in diet. . Comparisons were made using linear regression models comparing treatment, in subsets of the data by anatomical location. Only significant comparisons (p < 0.05) are listed. Bacterial richness is visualized in Figure 5.

| Treatment compared to control diet only group | Change in bacterial richness (SVs) ± Standard Error (rounded) | T-value | P-value |
| --- | --- | --- | --- |
| **Jejunum contents** |  |  |  |
| Control + 2.5% DSS | 40 ± 11 | 3.790 | 0.000608 *** |
| 10% Broccoli sprouts | 23 ± 10 | 2.766 | 0.009207 ** |
| 10% Broccoli sprouts + 2.5% DSS | 64 ± 10 | 6.141 | 5.05e-07 *** |
| **Jejunum scraping** |  |  |  |
| Control + 2.5% DSS | 52 ± 17 | 3.104 | 0.00434 ** |
| 10% Broccoli sprouts | 98 ± 16 | 6.230 | 1.16e-06 *** |
| 10% Broccoli sprouts + 2.5% DSS | 100 ± 15 | 6.708 | 3.35e-07 *** |
| **Cecum contents** |  |  |  |
| Control + 2.5% DSS | -13 ± 15 | -0.821 | 0.418 |
| 10% Broccoli sprouts | 68 ± 13 | 5.120 | 1.54e-05 *** |
| 10% Broccoli sprouts + 2.5% DSS | 70 ± 13 | 5.225 | 1.54e-05 *** |
| **Colon contents** |  |  |  |
| Control + 2.5% DSS | -12 ± 9 | -1.316 | 0.197 |
| 10% Broccoli sprouts | 43 ± 9 | 4.525 | 6.67e-05 *** |
| 10% Broccoli sprouts + 2.5% DSS | 90 ± 10 | 9.172 | 7.74e-11 *** |
| **Colon scraping** |  |  |  |
| Control + 2.5% DSS | 14 ± 12 | 1.225 | 0.129 |
| 10% Broccoli sprouts | 60 ± 12 | 5.220 | 1.78e-05 *** |
| 10% Broccoli sprouts + 2.5% DSS | 89 ± 12 | 7.749 | 2.81e-08 *** |

#### Beta diversity

Mice in the Broccoli+DSS group displayed a bacterial community that was distinct from that of mice in the Control+DSS group (Figure 6A,B). Bacterial taxa were characterized based on presence/absence (unweighted Jaccard Similarity, permANOVA: F = 16.90, p < 0.001) and presence/abundance (weighted Bray-Curtis Similarity, permANOVA: F = 29.97 p < 0.001). Interestingly, the bacterial community of the Broccoli+DSS mice was similar to the community of the Broccoli mice, suggesting that broccoli sprouts strongly protected against DSS-induced changes to the bacterial community (Figure 6A,B).

**Figure 6.**
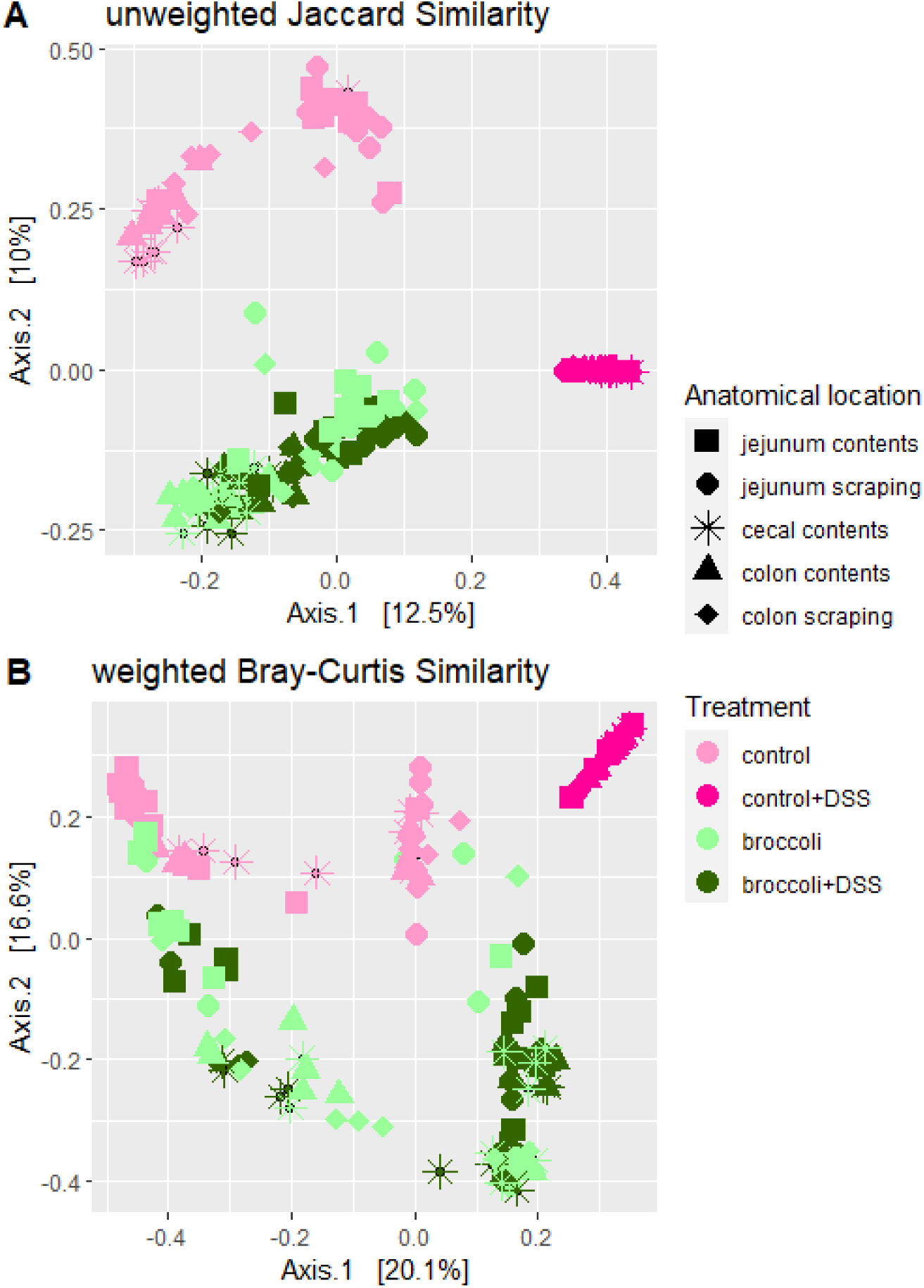
Principal coordinate analysis of bacterial community similarity along the intestines of mice under a DSS-induced model of chronic, relapsing colitis with or without 10% steamed broccoli sprouts in diet. Panel A was calculated with unweighted Jaccard Similarity to visualize differences in the taxonomic structure, and panel B with weighted Bray-Curtis to visualize structure and abundance.

#### Beta dispersion

Mice in the Broccoli+DSS group had a bacterial community dispersion (distance from centroids) larger than mice in the Control+DSS (Figure 6A,B), indicating more heterogeneity in the bacterial communities between mice fed broccoli sprouts. The Broccoli and Broccoli+DSS samples had a similar amount of dispersal between samples and the centroid for these treatments (beta dispersion, p > 0.05 for both comparisons, adjusted with TukeyHSD for multiple comparisons), indicating that broccoli sprouts negated any homogenization of the bacterial community caused by DSS. The Control and Control+DSS communities do not overlap and the Control+DSS community shows a tighter dispersal (beta dispersion, p < 0.05 for both comparisons, adjusted with Tukey’s HSD), indicating that DSS was a strong enough selective pressure to reduce the individual variation of bacterial communities within the group.

#### Biogeography

Mice with induced colitis on the broccoli sprouts diet (Broccoli+DSS) indicated clustering associated with each anatomical location (jejunum contents & scrapings, cecal contents, colon contents & scrapings) whereas mice without broccoli sprouts did not (Control+DSS, Figure 7A, permANOVA, uJS: F = 3.42, p < 0.001; wBC: F = 4.36, p < 0.001). The variation within the Control+DSS group was obscured by the strong comparisons to other groups in Figure 7A, so the Control+DSS samples were subset and demonstrated no biogeographic clustering (Figure 7B, permANOVA, p > 0.05, except marginally between jejunum scrapings vs. cecum contents, p = 0.04) when considering the bacterial community structure (unweighted Jaccard Similarity; Figure 7B) or structure and abundance (weighted Bray-Curtis, not shown).

**Figure 7.**
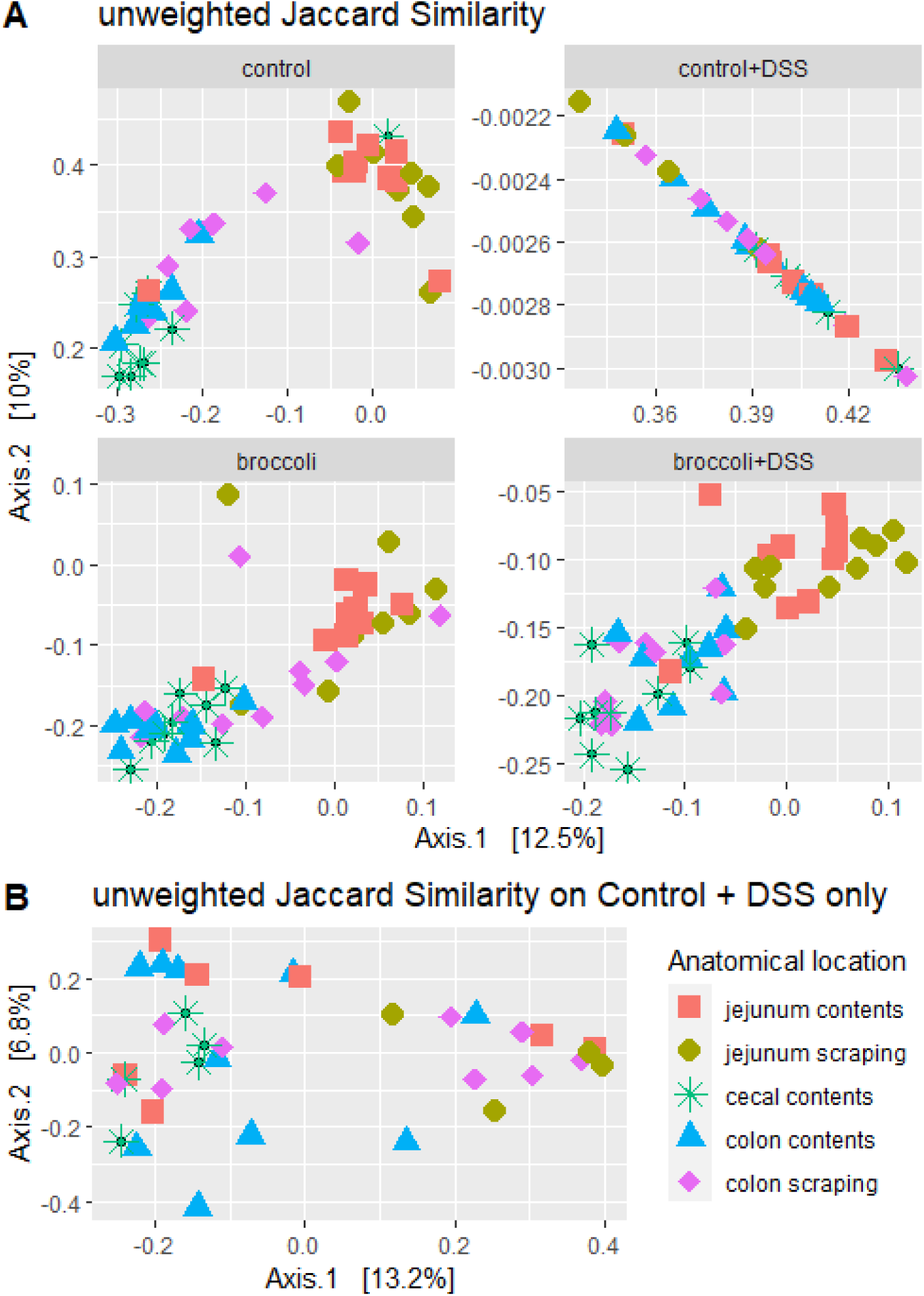
Principal coordinate analysis of bacterial communities from locations along the intestines of mice under a DSS-induced model of chronic, relapsing colitis with or without 10% steamed broccoli sprouts in diet: (A) All treatments (B) Control+DSS only. Bacterial communities in the jejunum, cecum, and colon were statistically different (permANOVA, p < 0.05) from each other in the treatment groups (A) Control, Broccoli, and Broccoli+DSS, were not statistically different from each other after multiple bouts of colitis in the (B) Control+DSS group.

#### Differential sequence variants

When identifying significant taxa which defined the Control samples by anatomical location, there were 188 significant (p < 0.05) taxa identified, with a permutational random forest model accuracy of 98% (Figure S2). Many of these defining taxa in the Control mice were species considered to be mouse commensals, such as *Dubosiella* spp. and strains from the Muribaculaceae and Lachnospiraceae families (Figure S2). The bacterial taxa which defined the Control+DSS mice included the *Clostridium sensu stricto* clade, and fewer commensal taxa were identified as important community members (Figure S3). The Broccoli sprout group was differentiated by *Dubosiella* spp., *Parasutterella excrementihominis*, and *Bacteroides* spp., among others (Figure S4). The Broccoli+DSS group had many of the same important taxa as the group consuming Broccoli sprouts without colitis (Figure S5), implying a strong selective pressure of the broccoli sprouts on the bacterial community structure. This was supported by the identification of ‘core taxa’ which were present in high abundance across at least 70% of the Broccoli and Broccoli+DSS samples (Figure S6A). Similar to the differential SV analysis, core bacteria in the Broccoli groups and core bacteria in the Control+DSS group did not share many similar taxa (Figure S6B). There were no bacterial SVs which were shared across at least 70% of samples in the Broccoli+DSS and the Control+DSS group (data not shown), indicating there was not a specific community which was enriched or associated with the DSS.

### Taxa with putative GLR metabolism capacity present in broccoli sprout fed mice even during colitis

#### Putative glucoraphanin metabolizers

Mice with induced colitis on the broccoli sprouts diet (Broccoli+DSS) indicated a dominant number of bacteria associated with the transformation of glucoraphanin into sulforaphane and distinct from mice not on a broccoli sprout diet which had relatively few (Control+DSS, Figure 8). The bacterial abundance in Broccoli+DSS mice was greatly reduced in the colon contents and somewhat reduced in the cecum contents and colon scrapings relative to the Broccoli mice. The bacterial abundance in Control+DSS mice increased across all anatomical sites relative to the Control mice. There were 309 bacterial SVs identified in samples to be associated with the expression of myrosinase-like enzymatic activity and the transformation of glucoraphanin to sulforaphane, which represented the genera *Bifidobacterium, Bacteroides, Enterococcus, Lactobacillus, Lactococcus, Pseudomonas, Staphylococcus,* and *Streptomyces* (21, 43, 44) 29, 54, 55). No *Faecalibacterium*, *Bacillus, Listeria, Pediococcus, Aerobacter, Citrobacter, Enterobacter, Escherichia, Salmonella, Paracolobactrum,* or *Proteus* were found in the samples (Figure 8). *Bacteroides* were predominant in both the Broccoli+DSS and Broccoli groups and much more abundant in comparison to all other genera across all samples (1,394,934 frequency across all samples compared to a total frequency of 6,791 for all other sequences; Figures 8,9).

**Figure 8.**
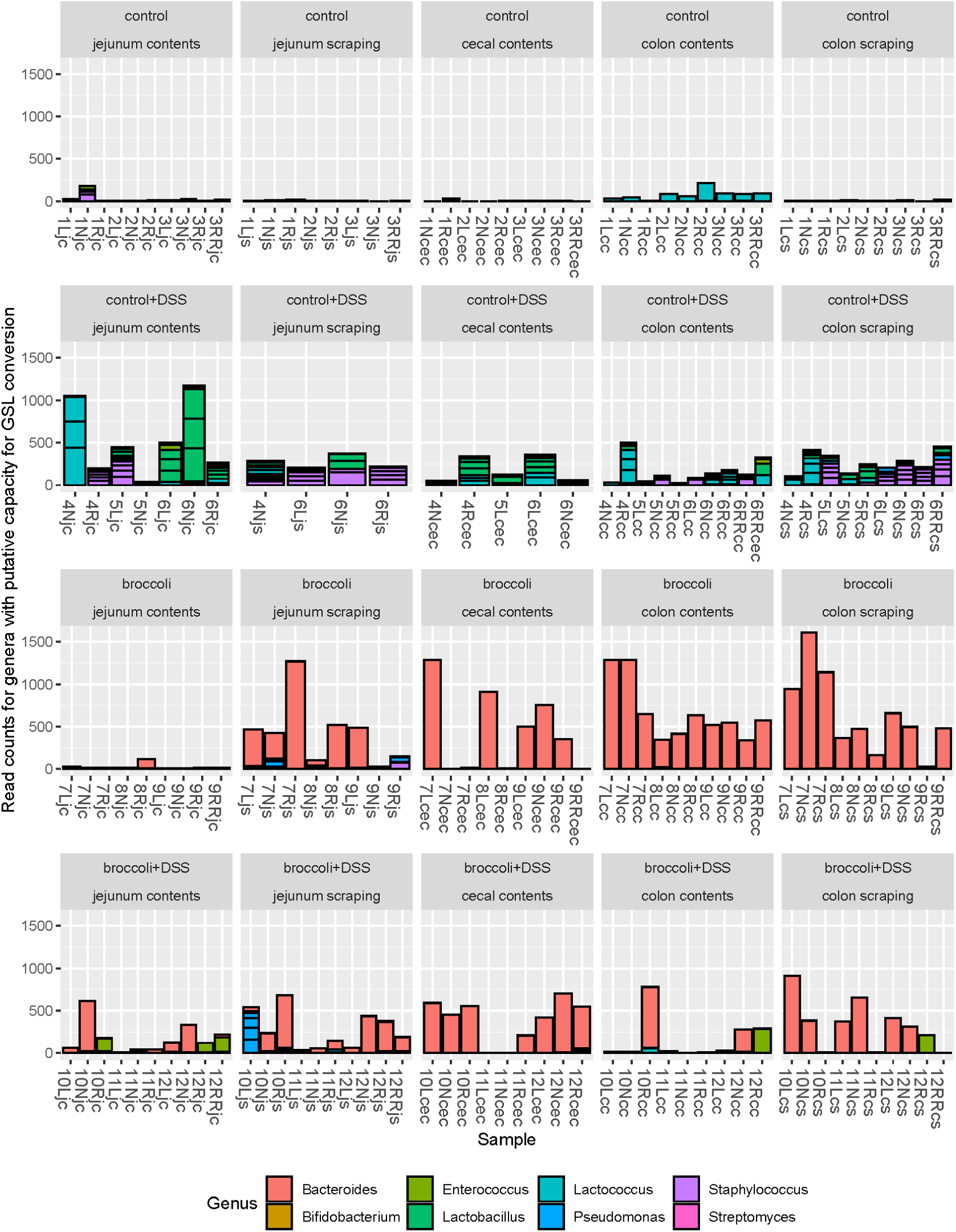
Bacterial sequence variants (SVs) belonging to the genera which have putative capacity to convert glucoraphanin to sulforaphane for mice under a DSS-induced model of chronic, relapsing colitis with or without 10% steamed broccoli sprouts in diet. Strains of bacteria in these genera have been demonstrated to perform myrosinase-like activity in the digestive tract, as reviewed in (43). Four treatment groups were used in a 34-day chronic, relapsing model of colitis: control diet, control diet with DSS added to drinking water, control diet adjusted with 10% by weight steamed broccoli sprouts, and 10% broccoli sprout diet with DSS added to drinking water.

**Figure 9.**
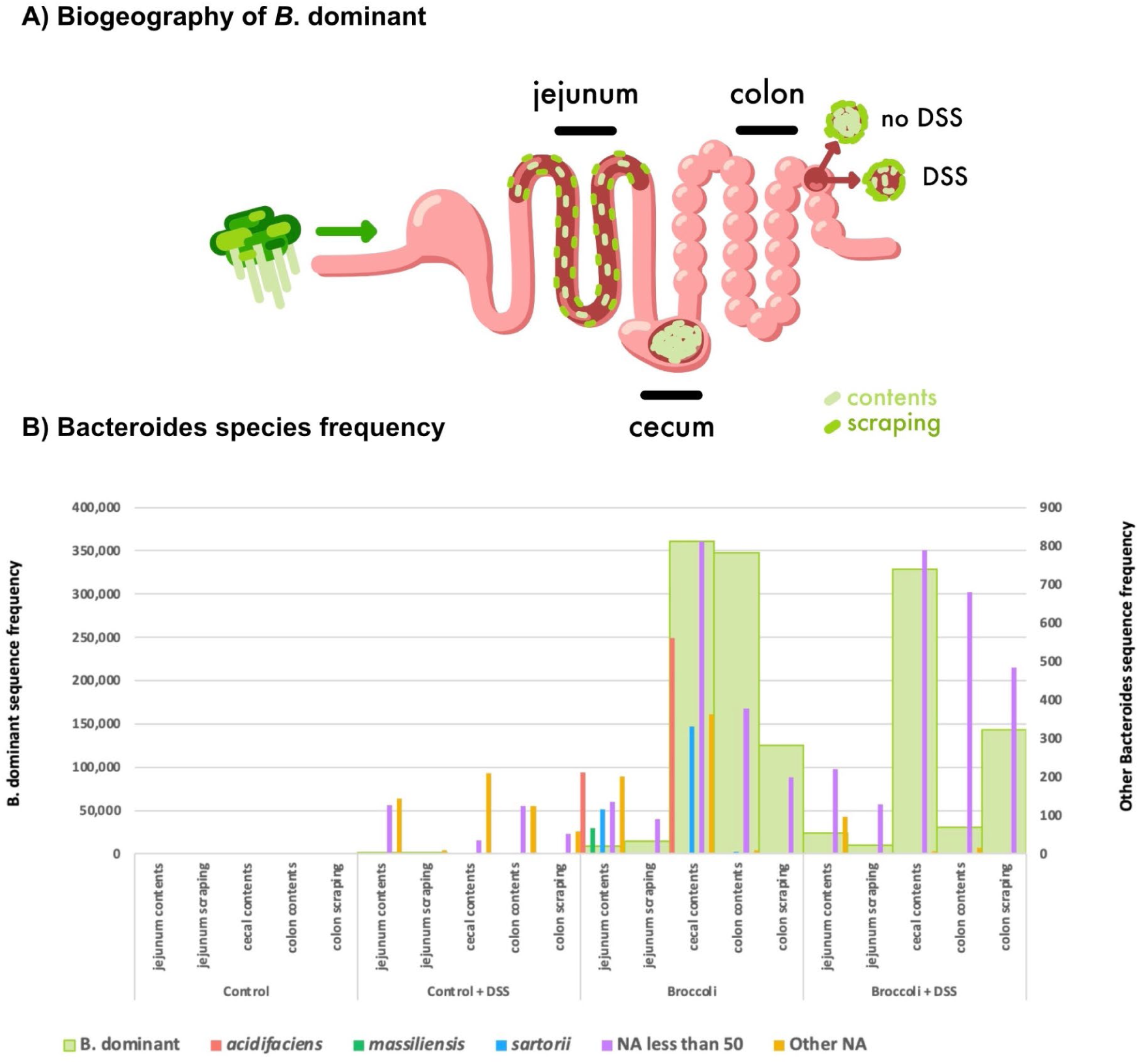
Biogeography of the dominant *Bacteroides* SV (*B.*-dominant) in the gut (A) and abundance of reads belonging to *Bacteroides* species by diet treatment and anatomical location in the gastrointestinal tract of mice. (A) *B.*-dominant was present in the jejunum contents and scrapings and abundant in the cecum contents, colon contents, and colon scrapings. DSS reduced *B*.-dominant colon content density. (B) The Silva Database identified *Bacteroides* species *acidifaciens*, *massiliensis*, and *sartorii*. Of the BLASTN identified species for *B.*-dominant SVs, only *B. thetaiotaomicron* was associated with broccoli-sprout-fed mice.

#### Bacteroides species

Several *Bacteroides* species were identified (*acidifaciens*, *massiliensis*, *sartorii*), but the dominant sequence variant was not identified to the species-level by the Silva Database taxonomy assignment (Figure 9A,B). This unidentified *Bacteroides* dominant SV (*B.-*dominant) was associated with multiple species in the NCBI Database (BLASTN): *Bacteroides thetaiotaomicron (B. theta)*, *Bacteroides faecis,* and *Bacteroides zhangwenhongii,* all with 100% Query Cover and Percent Identity (Figure 9B)*. Bacteroides theta* has been linked to myrosinase-like enzyme activity (21), but *Bacteroides faecis* and *Bacteroides zhangwenhongii* have not.

#### Bacteroides thetaiotaomicron (Bt VPI-5482)

Amplification of bacterial genes via qPCR indicated the presence of the *B. thetaiotaomicron* operon BT2159-BT2156, responsible for glucoraphanin metabolism, along the digestive tract (Figure S7). In each of the anatomical sites, BT2158 expression was most common followed by BT2157 and BT2156, BT2159, and then the regulatory gene BT2160 (Figure S7). Copy numbers were not significantly different for any individual gene by diet treatment and/or anatomical location (ANOVA, p > 0.05, adjusted for multiple comparisons with TukeyHSD). However, the operon’s GSL-metabolizing capacity is highest when all four genes are present (21), thus we also assessed the presence of the operon exhibited as collective gene copies, and samples demonstrated either high abundance (>100,000) or low abundance (<100; Figure 10).

**Figure 10:**
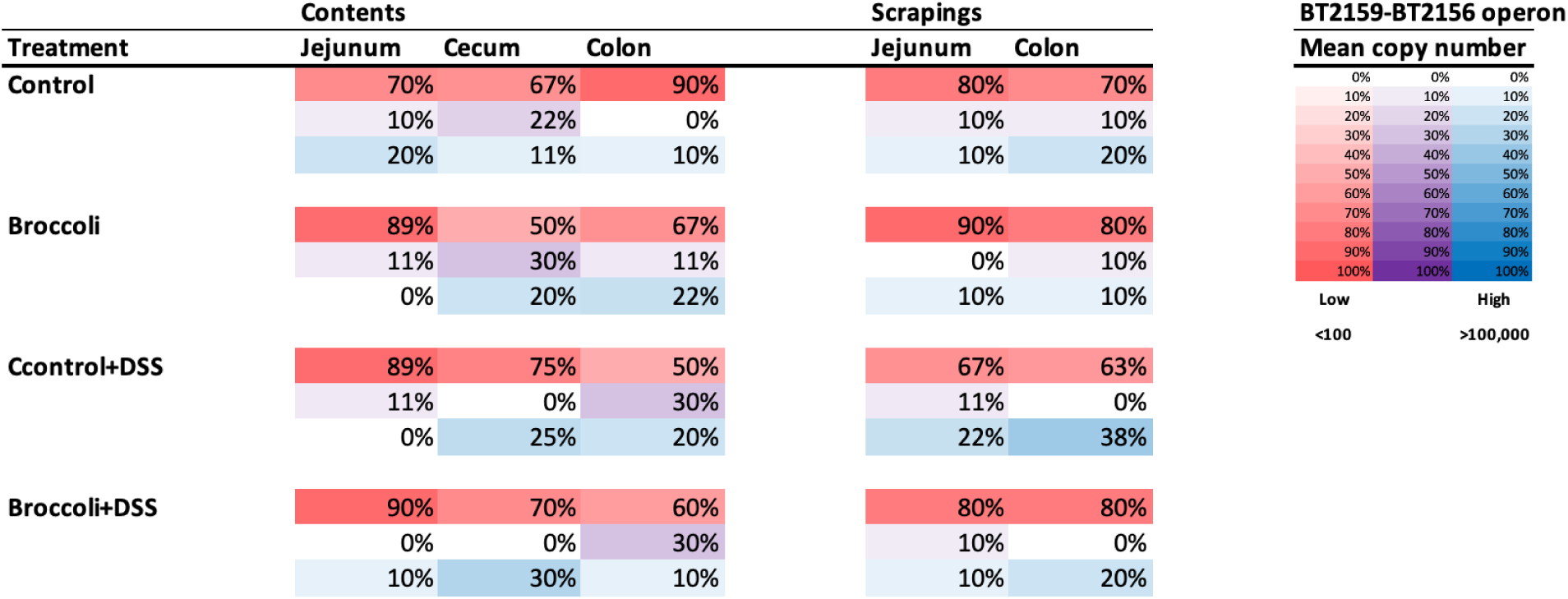
Prevalence of the operon BT2159-BT2156 in *Bacteroides thetaiotaomicron* (VPI-5482) in mice with and without chronic, relapsing DSS-induced colitis where diet was manipulated. Percentages are based on the number of samples (by diet and anatomical location) in each category of gene abundance (minimum mean copy numbers: red low <100, purple medium 100-100,000, blue high >100,000) across all 5 genes in the operon.

In the jejunum contents, 20% of Control mice had high presence of the operon BT2159-2156 (Figure 10). By contrast, the Broccoli group had low presence of the operon, which was associated with lower relative Bacteroidota (Figure S1), and the enrichment of SVs in the phyla Firmicutes, Actinobacteriota, and Proteobacteria. This had resulted in higher ratios of average Firmicutes/Bacteroidota (F/B: 72 v. 16), Actinobacteriota/Bacteroidota (A/B: 0.3 v. 0.1), and Proteobacteria/Bacteroidota (P/B: 0.6 v. 0.3). Operon presence in the Control+DSS and Broccoli+DSS jejunum contents was low. In the jejunum scrapings, more Control mice had a low presence of the operon, while more Broccoli mice exhibited higher presence (Figure 10). In the Control+DSS and Broccoli+DSS mice, the jejunum scrapings exhibited higher operon presence relative to the jejunum contents (Figure 10).

In the cecum contents, the presence of all 5 genes in the Broccoli group shows an increase in gene presence as compared to the jejunum samples in this group. Control+DSS and Broccoli+DSS groups had higher presence of the operon than in the groups without DSS. The operon presence in the Broccoli+DSS group is associated with *B.*-dominant and *Bacteroides* SVs (Figure 9), but the Control+DSS increase is associated with large-scale changes to the bacterial community (Figure S1): large reductions in Bacteroidota and Firmicutes, a large F/B ratio (118.5 v. 6.8), increased P/B (1.5 v. ∼0), and a significant increase in Actinobacteriota (A/B = 10.16 v. 0.01). The cecum samples on a broccoli diet were associated with higher counts of Bacteroidetes, Firmicutes, Actinobacteriota, and Proteobacteria and lower average ratios of F/B (10 v. 59), A/B (0.068 v. 4.8), and P/B (0.31 v. 0.70) in the cecum compared to groups on the control diet.

The colon contents in the Control mice showed limited operon presence, with higher abundance in scrapings, while the Control+DSS mice had higher operon presence in both the colon sites (Figure 10). The Broccoli mice showed higher operon presence in the contents as compared to the scrapings, while the Broccoli+DSS group had the opposite pattern.

## Discussion

### Broccoli sprouts protected mice against DSS-induced colitis

The anti-inflammatory effects of broccoli and broccoli sprout bioactives, in particular sulforaphane (SFN), have been well-established in various cell, animal, and human trials of IBD (17). However, studies have not elucidated the role of the whole gut communities, or patterns of biogeography, on the capacity of bacteria to metabolize the glucoraphanin (GLR) precursor to SFN. Juvenile SPF C57BL/6 mice (7-11 weeks) were fed a diet featuring 10% steamed broccoli sprouts, which was balanced by macronutrients and fiber, and which contained glucoraphanin but no plant myrosinase, thus gut microbiota were solely responsible for any production of the anti inflammatory sulforaphane. Under chronic, relapsing DSS-induced colitis (3 cycle regime), the steamed broccoli sprout diet group demonstrated mitigated weight loss; reduced Disease Activity Index, plasma lipocalin, and proinflammatory cytokines; the presence of bacterial taxa which are known to metabolize GLR to SFN; and retained biogeographical patterns of bacterial communities in different locations in the gut. Moreover, broccoli sprouts reduced the occurrence of fecal blood, which is known to increase the abundance of pathobionts and reduce commensals (45, 46).

Lipocalin is a neutrophil protein produced by epithelial cells and secreted luminally and through intestinal tissue where it is absorbed into the blood (47), and serves as a surrogate marker for intestinal inflammation in IBD (47). Plasma lipocalin was high in our Control+DSS mice, but low for the Broccoli+DSS diet which suggests that broccoli sprouts prevented severe physical destruction of the epithelium. Similarly, the lower concentration of plasma cytokines in the Broccoli+DSS mice indicates a lower innate immune response, and which may be the result of an intact epithelial barrier that prevented bacterial translocation from the lumen to intestinal tissues (34).

Broccoli sprout feeding was initiated before the introduction of DSS. This was done partly to acclimate mice to the diet, but primarily to investigate the efficacy of broccoli sprouts as a prevention strategy before initiating colitis. Previous studies have demonstrated its usefulness in alleviating symptoms during active disease states (13). To validate the feasibility of broccoli as a treatment for DSS-induced colitis, a 10% steamed broccoli sprout diet was used with male mice, as they are known to suffer more histopathological damage from DSS-induced colitis (48, 49). Regular access to fiber is critical to recruiting and retaining beneficial microbiota in the gut (50), and regular exposure to glucosinolates is needed to stimulate gut microbial conversion of GLR to SFN (51). In individuals who sporadically consume broccoli or broccoli sprouts, or consume them over short periods of time, the cooking preparation and the activity/inactivity of plant myrosinase were often the determining factors in the production of SFN, if any (51). However, directly including SFN in the diet is not feasible due to its instability (24). This approach would also hinder the potential for other benefits and the overall enjoyment of consuming a food rich in phenols and fiber.

We acknowledge that a major limitation in this study was only using male mice, as it is known that hormones mediate both the gut microbiome and the type and intensity of DSS-induced colitis. Thus, any future research on the application of this diet in people will require diversity in the gender of participants.

### Bacterial communities are highly responsive to DSS and broccoli sprouts

The broccoli sprout diet increased bacterial richness, regardless of DSS treatment, and as the control and sprout diets were balanced for total fiber it is likely that fiber was not solely responsible for this effect. In particular, broccoli sprouts increased bacterial richness in the colon. We previously showed that the concentration of SFN is highest in the colon after feeding C57BL/6 mice steamed broccoli sprout diets which has reduced myrosinase concentrations, suggesting that primary hydrolysis of SFN to GLR by microbial communities occurs in the colon (32). The current study found effects on microbial richness from the broccoli sprout diet manifested most strongly in the colon, supporting our previous findings.

When identifying significant taxa which defined the samples in different treatments, the Control mice contained a high abundance of known mouse commensal taxa, including *Dubosiella* spp. and strains from the Muribaculaceae and Lachnospiraceae families. The Control+DSS mice contained much lower abundance of known mouse commensal taxa and higher abundance of the *Clostridium sensu stricto* clade (52), which contains known pathogens such as *Clostridium perfringens* (53), and well-known butyrate-producing symbionts, such as *Clostridium butyricum* (54).

The Broccoli sprout group was also high in commensal taxa such as *Dubosiella* spp., and had enrichment of *Bacteroides* spp., a genus known to contain glucosinolate-metabolizing bacteria and modified by cruciferous vegetable consumption (55). The broccoli sprout diet enriched for *Parasutterella excrementihominis* in the gut, which is commonly found in human and murine gut communities and is associated with carbohydrate intake (56). It is possible that other breakdown products of glucosinolates, such as aromatic carbohydrate compounds in broccoli sprouts (57), caused this enrichment (57). *Parasutterella excrementihominis* has also been associated with inflammation in the gut in observation-based studies; however in metabolic pathway-based studies it appears to be an important contributor to nitrate reduction in the gut and in reducing stress related inflammation (58). Nitrates are low in raw broccoli and other cruciferous vegetables, but this can be increased by freezing (59). Thus, our diet preparation – which included freezing and freeze-drying steps – may have selected for nitrate-reducers. Also, outside of the mucosa, SFN has an anti-bacterial effect (17), which may explain why putative pathogenic genera were lower in Broccoli groups as compared to the Control+DSS group.

Importantly, the Broccoli+DSS group had many of the same important taxa as the group consuming Broccoli sprouts without colitis, implying that the selective pressure of the broccoli sprouts on the bacterial community structure may be stronger than that of the DSS. The two treatments were so similar that many SVs were found in high abundance across at least 70% of the samples in those mice. There were no bacterial SVs shared across at least 70% of the Broccoli+DSS and the Control+DSS group, indicating there was not a specific community which was enriched or associated with the DSS.

The inclusion of DSS in drinking water demonstrably affects mice and gut bacterial communities, but it does not necessarily reduce bacterial richness. Control+DSS mice hosted more bacterial richness than Control mice in the jejunum, and similar richness in the colon, although the distinct ordination clustering between Control and Control+DSS samples indicates different communities present regardless of similar richness, and the tight grouping of the Control+DSS community indicates a strong selective pressure. In part, Control+DSS mice may retain gut richness because certain bacteria, like *Proteus vulgaris,* can make use of DSS (60), while other novel commensals, like some *Bacteroides*, appear to offer protective effects (61). We did not identify any bacterial SVs belonging to *Proteus*, although we found significantly higher amounts of *Bacteroides* sp. in broccoli sprout-fed mice. *Bacteroides thetaiotaomicron* has been demonstrated to metabolize glucoraphanin to sulforaphane and offer protection against colitis in mice (21).

Across multiple mouse lineages, individualized bacterial communities responded differently to the inclusion of DSS, and the presence of *Duncaniella muricolitica* and *Alistipes okayasuensis* were implicated in better mouse health outcomes (62). Modeling of the gut microbiome indicates that certain taxa may act as keystone species – critical to building the rest of the community and particularly in circumstances in which the microbial community has been destabilized (63). However, no single important bacterial taxa has been identified in meta analyses of IBD in humans (64, 65), and some studies have pointed to the switch of bacterial commensals to acting as pathobionts as important (66).

There is considerable evidence that microbes provide metabolism of biologically inert glucosinolates to biologically active isothiocyanates, and several cultured bacterial strains from fermented foods and the digestive tract have been shown to perform this conversion (21, 43, 44). Additionally, a single bacterial equivalent of the plant myrosinase enzyme has not been identified, and there is the possibility that several enzymes in bacteria may work in concert to achieve metabolism of GSLs. Further, research is lacking to determine if the effect on the gut microbiome is metabolite (isothiocyanates)-mediated or precursor (glucosinolates)-mediated. The scope of research in this field should be expanded by investigating the concepts of biogeographic specificity of both bioactive production and absorption and microbial community dynamics.

### Biogeography reveals location-specific trends in bacterial communities

It has been well-demonstrated that distinct bacterial communities exist in different locations along the gastrointestinal tract in mammals, related to the local anatomical, environmental, nutritional, and host immunological conditions in different organs (67–69). Further, the effect of foods on the gut microbiome can be specific to an individual’s communities (27). Given the complexity of microbial community function, as well as the spatially explicit biochemical digestion activities of the host, it follows that the location of glucoraphanin metabolism may influence how well the host can absorb SFN, whether it may be distributed systematically, and where it will be effective at preventing or treating symptoms. It has previously been demonstrated in mouse trials using 2,4,6 trinitrobenzene sulfonic acid to chemically induce colitis that the mucosal-associated bacterial population is more affected by colitis than the lumen associated (digesta) community (36).

Here, we showed that DSS-induced colitis effectively erased the biogeographical specificity of communities in the mouse gastrointestinal tract, with the exception of communities in jejunum scrapings remaining distinct from those in cecal contents. Given the highly specific function of the cecum in separating fibers by size and furthering microbial fermentation, it is not surprising that it would be distinct. We sampled the jejunum because it is not often reported on in IBD studies, and does not always show effects of inflammation. We confirmed that bacteria in the jejunum are negatively affected by DSS. The DSS may be a stronger selective force than anatomical location in driving bacterial communities in the gut (61). For example, its destruction of the epithelial cell surface may change the microarchitecture and alter microbial attachment in the gut (70), and DSS increases the hydrophobicity of bile acids (71), which may affect microbial survival and ability to attach to host epithelial cells, thus increasing microbial washout through the gut. Any of these could possibly explain our findings that DSS eliminated biogeographical specificity of communities in the mouse gastrointestinal tract. However, more research would be needed to determine causation.

Mouse studies suggest biotransformation of glucoraphanin to SFN occurs in the colon (21, 25, 26), and we previously confirmed the majority of SFN accumulation takes place in the colon, with greater resolution using multiple locations along the GI tract (13). Critically, the cecum had low SFN, indicating that it is not responsible for hosting bacterial biotransformation of these bioactives (13). Here, we also showed that only a few bacterial taxa were estimated to be sourced in the cecum and creating sink populations in the colon, and of these, none were identified to be putatively responsible for metabolism of glucoraphanin. Collectively, this confirms that this is a valid model for generalization to the human gut.

### Putative GLR-metabolizers more abundant in sprout diets, but gene copies were not uniformly increased

Comparative analysis of the BLAST identification of SVs and qPCR of the gene copies in the operon BT2160-2156 suggest that the dominant Bacteroides SV identified by 16S rRNA sequencing largely represents *B. thetaiotaomicron*. The Broccoli diet (no DSS) induced the presence of *B.-*dominant in the jejunum contents and scrapings, and more so in the cecum contents and the colon contents and scrapings. The addition of DSS (Broccoli+DSS) noticeably reduced *B.-* dominant in the colon contents although it was maintained in the cecum and colon scrapings. Importantly, the results presented here used only DNA from viable cells, but not RNA, and it is known that *Bacteroides* and particularly *B. theta* exhibit highly situational carbohydrate and glycan digestion. Thus, the presence of *B. theta* or other putative GLR-metabolizers does not confirm their activity, which may explain why these taxa and GLR-metabolizing genes were also present in the Control and Control+DSS in low abundance.

Whereas the lumen (contents) may represent a transient view of a gut bacterial community, the mucosal-associated populations (scrapings) are more resistant to short-term changes to the diet or gut ecosystem. There was no *B.*-dominant present in the Control group’s jejunum contents or scrapings after 34 days of trial, but there were genes copies of the *B. theta* GLR-metabolizing operon, which suggests that gene copies were from the many unclassified Bacteroidota or *Bacteroides* SVs in the jejunum contents and scrapings sequencing samples, or from other genera containing an operon orthologous to BT2159-BT2156. By contrast, the Broccoli groups’s jejunum contents and scrapings include *B.*-dominant, with considerably higher frequencies in the scrapings, as well as higher prevalence of all 5 genes in the operon. This is further demonstrated in the DSS groups for both diets, in which the jejunum scrapings demonstrated higher prevalence of all 5 genes in the operon relative to the jejunum contents. Interestingly, the Control+DSS group had twice the prevalence of high copy numbers across all 5 genes, but no B.-dominant SVs, again suggesting the existence of the genes was due to other unclassified Bacteroidota or *Bacteroides*.

As chime flows from the jejunum to the cecum, it enters the less hostile (neutral to slightly alkaline) and primary fermentation environment of the cecum, where the bacterial density typically increases by orders of magnitude in healthy animals. In our samples, higher abundance of the operon was found in the cecum, as GLR-metabolizers are possibly benefiting from its slower transit time and more favorable fermentation environment. The presence of all 5 genes in the Broccoli group shows a dramatic increase in gene presence, as well as *B.*-dominant and *Bacteroides* SVs, as compared to the jejunum samples in this group. Interestingly, in the DSS groups for both diets, the prevalence of gene copies in the operon is higher than in the groups without DSS. The increase in the Broccoli+DSS group is associated with increased *B.*-dominant and *Bacteroides* SVs, but the Control+DSS increase is associated with phylum-scale changes to the bacterial community.

The Control and Control+DSS colon contents contained limited operon presence, with only slightly higher abundance in scrapings, and are largely absent of *B.*-dominant. The Broccoli and Broccoli+DSS groups contained more operon in the colon contents than the scrapings. In the colon contents in both broccoli sprout groups, *B.*-dominant was identified in a few mice, but *Bacteroides* counts were much higher and were not impacted by the DSS treatment. In the colon scrapings in both broccoli sprout groups, *B.*-dominant was more prevalent and dominant, suggesting the possibility of *B*.-dominant GLR-metabolism in the mucosal-associated sites, and other *Bacteroides* providing GLR-metabolism in the Broccoli+DSS colon contents.

### Considerations for the application of this work

Access to fresh or frozen broccoli and broccoli sprouts, the cooking preparation, and the ability to regularly consume these vegetables will have implications for the feasibility and success of a dietary intervention for preventing or reducing inflammation in the gut. For example, sprouts from various plants have been implicated in food-borne illness because of their proximity to soil, and people may be wary of or discouraged from consuming raw or slightly cooked sprouts. However, more recent research has shown that sprouts can be consumed safely, especially with improvements in hygiene and agricultural regulations, as well as in food processing (72). Further, broccoli sprouts can be grown at home in windowsill seed-bed germinators without requiring soil, gardening tools, or specialized gardening knowledge, which could ease the financial burden of purchasing healthy foods (73). This may prove to be particularly important in areas without access to healthcare or affordable prescriptions, or in areas without close proximity to fresh, healthy fruits and vegetables (5, 74, 75) as this can preclude being able to make the dietary recommendations set forth by medical professionals (76). Including broccoli sprouts as 10% of the diet could be potentially too high for IBD patients to comply with, and future studies on the application of this diet will require a deeper understanding of the biological, microbiological, immunological, as well as social and logistical factors involved in dietary interventions in people.

## Materials and Methods

### Diet preparation

Jonathan’s Sprouts™ (Rochester, Massachusetts, USA) broccoli sprouts were purchased from a grocery store (Bangor, Maine, USA) and steamed in a double boiler for 10 minutes, immediately cooled down, and stored in a -80℃ freezer until freeze-drying (University of Maine Food Pilot Plant, Orono, Maine, USA). The freeze-dried broccoli sprouts were ground into a fine powder and mixed with purified AIN93G rodent base diet (Envigo, now Inotiv, Indianapolis, Indiana, USA) to a concentration of 10% by weight. The AIN93G diet contains 50 g/kg of fiber as cellulose (77), and broccoli sprouts contain 35 g/kg fiber, thus the broccoli sprout-fed mice consumed a similar amount of fiber on a per weight basis. The diets were measured to balance their weight prior to feedings; however, we did not measure food intake or refusals. Anecdotal animal care notes and visual observation of cages across multiple previous studies indicate that control and broccoli sprout groups do not consume different amounts of food (Li and Ishaq, personal communications.

Our previous work assessed the effects of different diet preparations and the percentage of broccoli sprouts, and found that 5-10% broccoli sprouts by weight reliably produces consistent anti-inflammatory results in mice (13). For this study, we chose to use 10% steamed broccoli sprouts both to assess the microbial conversion of GLR to SFN, and to ensure that the intervention would have a strong effect (48, 49, 78). Diet pellets were formed using a silicone mold to ensure consistent sizing, and allowed to dry at room temperature for up to 48 hours in a chemical safety hood to facilitate moisture evaporation, and after drying were stored in sealable plastic bags in a 20℃ freezer.

### DSS colitis model

The dextran sodium sulfate (DSS) mouse model is widely used to study human Ulcerative Colitis (UC) (41, 79), and has been used for studying diet-sourced anti-inflammatories for the prevention of IBD (40, 41). Administration of DSS in drinking water modifies the expression of tight junction proteins in intestinal epithelial cells, leading to a leaky epithelial barrier (80). This is followed by goblet cell depletion, erosion, ulceration, and infiltration of neutrophils into the lamina propria and submucosa (81), triggering the innate immune response (82, 83).

Forty male, 6-week-old, specific pathogen free (SPF) C57BL/6 mice (*Mus musculus*) were purchased from the Jackson Laboratory (Bar Harbor, Maine, U.S.) and transferred to the animal facility at the University of Maine (Orono, Maine, U.S.). The animal protocol (IACUC protocol A2020-01-04) was approved by the University Committee on the Use and Care of Animals, and all biosafety work was approved by the Institutional Biosafety Committee (protocol SI-092220). The mice were acclimated to the facility for 7 days (day -6 to 0), during which they received *ad libitum* autoclaved tap water and the AIN-93G purified rodent diet (control diet). After initial acclimation, the mice were randomly assigned to one of 4 experimental groups beginning on experimental day 1: control diet without DSS treatment (Control), 10% steamed broccoli sprout diet without DSS treatment (Broccoli), control diet with DSS treatment (Control+DSS), and 10% steamed broccoli sprout diet with DSS treatment (Broccoli+DSS). All experimental groups were on 7 days of their respective diets (control or 10% steamed broccoli sprout), after which DSS (Alfa Aesar, molecular weight ∼40 kD (39)) was added to the drinking water of the DSS treatment groups to a final concentration of 2.5%. Mice were given DSS for 5 days, followed by a recovery period for 5 - 7 days. This was repeated for a total of 3 cycles to induce chronic colitis (40, 84). Mice were sacrificed and tissue collected after the third round of DSS, on day 35 of the experiment.

Bodyweight, fecal blood, and fecal consistency were used to calculate Disease Activity Index (DAI) scores (83). Fecal samples were collected every 2 - 3 days throughout the trial, and daily during the DSS cycles. Body weights and DAI were analyzed using 2-way ANOVA generated with R to compare differences between treatments for each day. A generalized additive model (GAM) was used in R to compare DAI differences (R-sq.(adj) = 0.861, Deviance explained = 86.4%, GCV = 0.036031) by treatment across the entire study, using Mouse ID to account for repeated measures.

Lipocalin-2 concentration in the plasma samples were determined by a mouse Lipocalin 2/NGAL DuoSet ELISA kit (R & D Biosystems, USA) following the manufacturer’s instructions. Lipocalin is a neutrophil protein that binds bacterial siderophores and serves as a surrogate marker for intestinal inflammation in IBD (47). The readings at wavelengths of 540 nm and 450 nm were measured by a Thermo Scientific Varioskan LUX Multimode Microplate Reader. The readings at 540 nm were subtracted from the readings at 450 nm to correct for the discoloration of the solution. ANOVA was used to compare lipocalin values.

We analyzed several cytokines in the mouse plasma samples which were collected at the end of the study. CCL4, also known as macrophage inflammatory protein 1β (MIP1β), is a chemokine that plays a critical role in regulating the migration and activation of immune cells during inflammatory processes (42). Interleukins IL-1β and IL-6; and tumor necrosis factor alpha (TNF-α) are pro-inflammatory cytokines that play key roles in the regulation of the immune response (85–87). The concentrations of mouse CCL4/MIP-1β, IL-1β/IL1F2, IL6, and TNF-a were analyzed using the Simple Plex Ella Automated Immunoassay System (Ella) from R&D Biosystems, USA. Mouse serum samples were diluted 10-fold using the reagent diluent provided with the kit (SPCKA-MP-007374, Protein Simple, Bio-Techne), and the concentrations were determined following the manufacturer’s instructions. The Ella system was used to perform the immunoassay, and mean values were calculated for each analyte. The resulting data were used for statistical analyses using ANOVA.

After euthanasia, lumen-associated (digesta contents) and mucosal-associated (epithelial scrapings) microbial community samples were collected from the jejunum, cecum (contents only), and colon for DNA extraction, gene quantification, and community sequencing as described below.

### Bacterial community sequencing and analysis

Immediately following euthanasia of the mice, digesta (lumen contents) and epithelial associated (tissue scrapings) microbial community samples were collected from the jejunum, the cecum (contents only), and along the entire colon, as the inflamed mice had short colons and limited amounts of digesta present. All samples of gut microbiota were gently homogenized with vortexing, then treated with propidium monoazide (PMA; BioTium) following kit protocols at a final concentration of 25 μm. PMA covalently binds to relic/free DNA and DNA inside compromised/dead cell membranes, and prevents amplification in downstream protocols to preclude dead DNA from the sequence data (88).

Following PMA treatment, bulk DNA was extracted from bacterial communities (n = 200 samples), or no-template (water) control samples (n = 10, one for each extraction batch) using commercially available kits optimized for water and tissue-based microbial communities (Quick DNA Fecal/Soil Kit, Zymo Research). DNA extract was roughly quantified and purity-checked with a Thermo Scientific™ NanoDrop™ OneC Microvolume UV-Vis Spectrophotometer (Thermo Scientific, Waltham, MA, U.S.). Samples underwent DNA amplicon sequencing of the 16S rRNA gene V3-V4 region, using primers 341F and 806R and protocols consistent with The Earth Microbiome Project (89), and sequenced on an Illumina MiSeq platform using the 2 x 300 nt V3 kit (Molecular Research Labs, Clearwater, TX, U.S.). Raw sequence data (fastq files and metadata) is available from NCBI through BioProject Accession number PRJNA911821.

Amplicon sequence data was processed using previously curated workflows in the Ishaq Lab (R code supplied as Supplemental Material), which used the DADA2 pipeline ver. 1.26 (90) in the R software environment ver. 4.1.1 (91). The dataset started with 46,581,832 paired raw reads, and based on initial quality assessment only the forward reads were processed. Trimming parameters were designated based on visual assessment of the aggregated quality scores at each base from all samples (plotQualityProfile in DADA2): the first 10 bases were trimmed, sequences were trimmed to 225 bases in length, and were discarded if they had ambiguous bases, more than two errors, or matched the PhiX version 3 positive control (Illumina; FC-110-3001). After filtering, 34,009,802 non-unique forward/read 1 sequences remained.

The DADA algorithm was used to estimate the error rates for the sequencing run, dereplicate the reads, pick sequence variants (SVs) which represent ‘microbial individuals’, and remove chimeric artifacts from the sequence table. Taxonomy was assigned using the Silva taxonomic training data version 138.1 (92) down to species where possible, and reads matching chloroplasts and mitochondria taxa were removed using the dplyr package (93). No-template control samples were used to remove contaminating sequences from the samples by extraction batch (94). The sequence table, taxonomy, and metadata were combined for each experiment using the phyloseq package (95), which was also used for basic visualization and statistical analysis in combination with other packages. Samples from one mouse (4L, in the Control+DSS group) were dropped from further analysis as they were outliers on all visualizations and may have been contaminated during DNA extraction.

Normality was checked using a Shapiro-Wilkes test on alpha diversity metrics generated from rarefied data, including observed richness, evenness, and Shannon diversity. Linear models were run for comparisons of alpha diversity metrics to compare by sample type, (lme4 package (96)), in which anatomical location and diet treatment were used as fixed effects, and mouse ID used to control for repeated sampling as needed. Generalized additive models were used to assess trends in alpha diversity using time as a smoother (97). Jaccard unweighted similarity was used to calculate sample similarity based on community membership (species presence/absence), visualized with non-parametric multidimensional scaling, and tested with permutational analysis of variance (permANOVA) by using the vegan package (98). Random forest feature prediction with permutation was used to identify differentially abundant SVs based on factorial conditions (99). Plots were made using the ggplot2 (100), ggpubr (101), and phyloseq packages.

Source Tracker algorithms which had been modified for the R platform (102, 103) were used to identify source:sink effects based on anatomical location. This was used to determine if the cecum could be the source for population sinks in the colon, as a proxy for the model’s applicability to the human gut anatomical features and microbial communities. A total of 142 SVs were identified as possibly sourced from the cecum, and 95 SVs were estimated to make up >1% of the proportion of sources (Figure S8). The putative GLR converting bacteria in the Broccoli and Broccoli+DSS mouse gut samples were not among those taxa identified as sourced in the cecum.

### Quantitative PCR for glucoraphanin metabolism genes

DNA extract from all gut locations was used to assess microbial potential for glucoraphanin conversion. Bacterial genes, gene sequences, and primers associated with conversion of GSLs to ITCs identified in *Bacteroides thetaiotaomicron* VPI-5482 (21) were used, and are listed in Table S1. These primers were evaluated for validity using IDT’s OligoAnalyzer (www.idtdna.com) and NCBI primer blast (https://www.ncbi.nlm.nih.gov/tools/primer-blast/). The identified genes from *Bacteroides thetaiotaomicron* VPI-5482 were analyzed by using the ApE-A plasmid Editor v3.1.3 to optimize and validate the primers for optimal GC content and melting temperature.

Quantitative polymerase chain reaction (qPCR) was performed on extracted DNA using the Applied Biosystems QuantStudio 6 Flex Real-Time PCR system (Applied Biosystems, Foster City, CA, USA). Luna® Universal qPCR Master Mix and primer sets BT2156-BT2160 were used to quantify copy numbers of glucoraphanin-metabolizing genes previously identified from *Bacteroides thetaiotaomicron*. Primers were diluted to 10 µM, for a final concentration of 0.25 µM in each well. Each primer set required one cycle of 50°C for 2 min, one cycle of 95°C for 1 min, and forty cycles of 95°C for 15 sec, 60°C for 30 sec and 72°C at either 20, 25 or 30 sec depending on the primer set. A standard curve for each gene was created using a set of 7 serially diluted geneblocks (IDT). All samples and standards were run in triplicate, across two 384-well plates with negative controls and standards included on both plates. Primer sequences, geneblock sequences for standards, and protocols are listed in Table S1. Sample gene copy numbers were calculated using the standard curve in the Quantstudio analysis software. Data were visualized and statistically analyzed in R using analysis of variance models corrected for multiple comparisons using Tukey’s HSD. Similar to the bacterial community data, sample 4L was dropped from the qPCR data analysis as an outlier/possible contamination.

## Acknowledgements

All authors have read and approved the final manuscript. The authors thank Jess Majors, University of Maine, for her kind and detailed care of the mice during the trial, and for Ellie Pelletier for her informal review of the manuscript. This project was supported by the USDA National Institute of Food and Agriculture through the Maine Agricultural & Forest Experiment Station: Hatch Project Numbers ME022102 and ME022329 (Ishaq) and ME022303 (Li) which supported Johanna Holman; the USDA-NIFA-AFRI Foundational Program [Li and Chen; USDA/NIFA 2018-67017-27520/2018-67017-36797]; and the National Institute of Health [Li and Ishaq; NIH/NIDDK 1R15DK133826-01] which supported Marissa Kinney, Timothy Hunt, and Benjamin Hunt. Lola Holcomb was supported by US National Science Foundation One Health and the Environment (OG&E): Convergence of Social and Biological Sciences NRT program grant DGE-1922560, and through the UMaine Graduate School of Biomedical Sciences and Engineering.

## Author Contributions

Conceptualization, S.L.I., Y.L., T.Z., G.C., G.M; Methodology, S.L.I., Y.L., T.Z., G.M., P.M., J.H.; Software, S.L.I.; Formal Analysis, J.H., M.K., S.L.I., T.H., B.H.; Investigation, J.H., L.C., J.B.; G.C., D.B.; Resources, S.L.I., Y.L., T.Z.; Data Curation, S.L.I., J.H.; Writing – Original Draft, J.H., L.C., S.L.I., Y.L.; Writing - Review and Editing; S.L.I., Y.L., T.Z., G.M., P.M., J.H., L.C., J.B., G.C., D.B., T.H., B.H., L.H. M.K.; Visualization, J.H., S.L.I., T.H., B.H.; Supervision, J.H., S.L.I., Y.L., T.Z.; Project Administration, S.L.I., Y.L., T.Z; Funding Acquisition, S.L.I., Y.L., T.Z., G.C.

## Supplemental Materials Legends

### Supplemental Figures and Table

Additional data visualizations are included to demonstrate minor points or null results.

### Supplemental Material

The R code that was used to clean, visualize, and statistically analyze the bacterial community sequencing data presented.

## References

1. Peery AF, Crockett SD, Murphy CC, Lund JL, Dellon ES, Williams JL, Jensen ET, Shaheen NJ, Barritt AS, Lieber SR, Kochar B, Barnes EL, Fan YC, Pate V, Galanko J, Baron TH, Sandler RS. 2019. Burden and Cost of Gastrointestinal, Liver, and Pancreatic Diseases in the United States: Update 2018. Gastroenterology 156:254–272.e11.

2. Singh S, Qian AS, Nguyen NH, Ho SKM, Luo J, Jairath V, Sandborn WJ, Ma C. 2022. Trends in U.S. Health Care Spending on Inflammatory Bowel Diseases, 1996-2016. Inflamm Bowel Dis 28:364–372.

3. Lu Q, Yang M-F, Liang Y-J, Xu J, Xu H-M, Nie Y-Q, Wang L-S, Yao J, Li D-F. 2022. Immunology of Inflammatory Bowel Disease: Molecular Mechanisms and Therapeutics. J Inflamm Res 15:1825–1844.

4. Rubin DC, Shaker A, Levin MS. 2012. Chronic intestinal inflammation: Inflammatory bowel disease and colitis-associated colon cancer. Frontiers in Immunology 3:00107.

5. Walker RE, Keane CR, Burke JG. 2010. Disparities and access to healthy food in the United States: A review of food deserts literature. Health Place 16:876–884.

6. Banerjee S, Ghosh S, Sinha K, Chowdhury S, Sil PC. 2019. Sulphur dioxide ameliorates colitis related pathophysiology and inflammation. Toxicology 412:63–78.

7. Lewis JD, Abreu MT. 2017. Diet as a Trigger or Therapy for Inflammatory Bowel Diseases. Gastroenterology 152:398–414.e6.

8. Armstrong HK, Bording-Jorgensen M, Santer DM, Zhang Z, Valcheva R, Rieger AM, Sung-Ho Kim J, Dijk SI, Mahmood R, Ogungbola O, Jovel J, Moreau F, Gorman H, Dickner R, Jerasi J, Mander IK, Lafleur D, Cheng C, Petrova A, Jeanson T-L, Mason A, Sergi CM, Levine A, Chadee K, Armstrong D, Rauscher S, Bernstein CN, Carroll MW, Huynh HQ, Walter J, Madsen KL, Dieleman LA, Wine E. 2022. Unfermented β-fructan Fibers Fuel Inflammation in Select Inflammatory Bowel Disease Patients. Gastroenterology https://doi.org/10.1053/j.gastro.2022.09.034.

9. Tse G, Eslick GD. 2014. Cruciferous vegetables and risk of colorectal neoplasms: a systematic review and meta-analysis. Nutr Cancer 66:128–139.

10. Li Y, Zhang T, Schwartz SJ, Sun D. 2011. Sulforaphane Potentiates the Efficacy of 17 Allylamino 17-Demethoxygeldanamycin Against Pancreatic Cancer Through Enhanced Abrogation of Hsp90 Chaperone Function. Nutrition and Cancer https://doi.org/10.1080/01635581.2011.596645.

11. Li Y, Zhang T, Korkaya H, Liu S, Lee H-F, Newman B, Yu Y, Clouthier SG, Schwartz SJ, Wicha MS, Sun D. 2010. Sulforaphane, a dietary component of broccoli/broccoli sprouts, inhibits breast cancer stem cells. Clin Cancer Res 16:2580–2590.

12. Angelino D, Jeffery E. 2014. Glucosinolate hydrolysis and bioavailability of resulting isothiocyanates: Focus on glucoraphanin. J Funct Foods 7:67–76.

13. Zhang T, Holman J, McKinstry D, Trindade BC, Eaton KA, Mendoza-Castrejon J, Ho S, Wells E, Yuan H, Wen B, Sun D, Chen GY, Li Y. 2022. A steamed broccoli sprout diet preparation that reduces colitis via the gut microbiota. J Nutr Biochem 112:109215.

14. Fahey JW, Stephenson KK, Wade KL, Talalay P. 2013. Urease from *Helicobacter pylori* is inactivated by sulforaphane and other isothiocyanates. Biochem Biophys Res Commun 435:1–7.

15. Fahey JW, Wehage SL, Holtzclaw WD, Kensler TW, Egner PA, Shapiro TA, Talalay P. 2012. Protection of humans by plant glucosinolates: efficiency of conversion of glucosinolates to isothiocyanates by the gastrointestinal microflora. Cancer Prev Res (Phila) 5:603–611.

16. Wei L-Y, Zhang J-K, Zheng L, Chen Y. 2022. The functional role of sulforaphane in intestinal inflammation: a review. Food Funct 13:514–529.

17. Holman J, Hurd M, Moses P, Mawe G, Zhang T, Ishaq SL, Li Y. 2023. Interplay of Broccoli/Broccoli Sprout Bioactives with Gut Microbiota in Reducing Inflammation in Inflammatory Bowel Diseases. J Nutr Biochem 113:109238.

18. Zhang Y, Tan L, Li C, Wu H, Ran D, Zhang Z. 2020. Sulforaphane alter the microbiota and mitigate colitis severity on mice ulcerative colitis induced by DSS. AMB Express 10:119.

19. Li Y, Wicha MS, Schwartz SJ, Sun D. 2011. Implications of cancer stem cell theory for cancer chemoprevention by natural dietary compounds. J Nutr Biochem 22:799–806.

20. Bricker GV, Riedl KM, Ralston RA, Tober KL, Oberyszyn TM, Schwartz SJ. 2014. Isothiocyanate metabolism, distribution, and interconversion in mice following consumption of thermally processed broccoli sprouts or purified sulforaphane. Mol Nutr Food Res 58:1991–2000.

21. Liou CS, Sirk SJ, Diaz CAC, Klein AP, Fischer CR, Higginbottom SK, Erez A, Donia MS, Sonnenburg JL, Sattely ES. 2020. A Metabolic Pathway for Activation of Dietary Glucosinolates by a Human Gut Symbiont. Cell 180:717–728.e19.

22. Elfoul L, Rabot S, Khelifa N, Quinsac A, Duguay A, Rimbault A. 2001. Formation of allyl isothiocyanate from sinigrin in the digestive tract of rats monoassociated with a human colonic strain of *Bacteroides thetaiotaomicron*. FEMS Microbiol Lett 197:99–103.

23. Shapiro TA, Fahey JW, Wade KL, Stephenson KK, Talalay P. 1998. Human metabolism and excretion of cancer chemoprotective glucosinolates and isothiocyanates of cruciferous vegetables. Cancer Epidemiol Biomarkers Prev 7:1091–1100.

24. Yagishita Y, Fahey JW, Dinkova-Kostova AT, Kensler TW. 2019. Broccoli or Sulforaphane: Is It the Source or Dose That Matters? Molecules 24:3593.

25. Li F, Hullar MAJ, Beresford SAA, Lampe JW. 2011. Variation of glucoraphanin metabolism in vivo and ex vivo by human gut bacteria. Br J Nutr 106:408–416.

26. Lai R-H, Miller MJ, Jeffery E. 2010. Glucoraphanin hydrolysis by microbiota in the rat cecum results in sulforaphane absorption. Food Funct 1:161–166.

27. Johnson AJ, Vangay P, Al-Ghalith GA, Hillmann BM, Ward TL, Shields-Cutler RR, Kim AD, Shmagel AK, Syed AN, Personalized Microbiome Class Students, Walter J, Menon R, Koecher K, Knights D. 2019. Daily Sampling Reveals Personalized Diet-Microbiome Associations in Humans. Cell Host Microbe 25:789–802.e5.

28. Amato KR, Arrieta M-C, Azad MB, Bailey MT, Broussard JL, Bruggeling CE, Claud EC, Costello EK, Davenport ER, Dutilh BE, Swain Ewald HA, Ewald P, Hanlon EC, Julion W, Keshavarzian A, Maurice CF, Miller GE, Preidis GA, Segurel L, Singer B, Subramanian S, Zhao L, Kuzawa CW. 2021. The human gut microbiome and health inequities. Proc Natl Acad Sci U S A 118:e2017947118.

29. Hillman ET, Lu H, Yao T, Nakatsu CH. 2017. Microbial Ecology along the Gastrointestinal Tract. Microbes Environ 32:300–313.

30. Lynch SV, Pedersen O. 2016. The Human Intestinal Microbiome in Health and Disease. N Engl J Med 375:2369–2379.

31. Pereira FC, Berry D. 2017. Microbial nutrient niches in the gut. Environ Microbiol 19:1366–1378.

32. Press AG, Hauptmann IA, Hauptmann L, Fuchs B, Fuchs M, Ewe K, Ramadori G. 1998. Gastrointestinal pH profiles in patients with inflammatory bowel disease. Aliment Pharmacol Ther 12:673–678.

33. Compare D, Pica L, Rocco A, De Giorgi F, Cuomo R, Sarnelli G, Romano M, Nardone G. 2011. Effects of long-term PPI treatment on producing bowel symptoms and SIBO. European Journal of Clinical Investigation https://doi.org/10.1111/j.1365-2362.2010.02419.x.

34. Swidsinski A, Ladhoff A, Pernthaler A, Swidsinski S, Loening-Baucke V, Ortner M, Weber J, Hoffmann U, Schreiber S, Dietel M, Lochs H. 2002. Mucosal flora in inflammatory bowel disease. Gastroenterology 122:44–54.

35. Gevers D, Kugathasan S, Denson LA, Vázquez-Baeza Y, Van Treuren W, Ren B, Schwager E, Knights D, Song SJ, Yassour M, Morgan XC, Kostic AD, Luo C, González A, McDonald D, Haberman Y, Walters T, Baker S, Rosh J, Stephens M, Heyman M, Markowitz J, Baldassano R, Griffiths A, Sylvester F, Mack D, Kim S, Crandall W, Hyams J, Huttenhower C, Knight R, Xavier RJ. 2014. The treatment-naive microbiome in new onset Crohn’s disease. Cell Host Microbe 15:382–392.

36. Kozik AJ, Nakatsu CH, Chun H, Jones-Hall YL. 2019. Comparison of the fecal, cecal, and mucus microbiome in male and female mice after TNBS-induced colitis. PLoS One 14:e0225079.

37. Xu P, Lv T, Dong S, Cui Z, Luo X, Jia B, Jeon CO, Zhang J. 2022. Association between intestinal microbiome and inflammatory bowel disease: Insights from bibliometric analysis. Comput Struct Biotechnol J 20:1716–1725.

38. Boger-May A, Reed T, LaTorre D, Ruley-Haase K, Hoffman H, English L, Roncagli C, Overstreet A-M, Boone D. 2022. Altered microbial biogeography in an innate model of colitis. Gut Microbes 14:2123677.

39. da Costa Goncalves F, Schneider N, Mello HF, Passos EP, da Rosa Paz AH. 2013. Characterization of Acute Murine Dextran Sodium Sulfate (DSS) Colitis: Severity of Inflammation is Dependent on the DSS Molecular Weight and Concentration. Acta Scientiae Veterinari 41:1–9.

40. Wirtz S, Popp V, Kindermann M, Gerlach K, Weigmann B, Fichtner-Feigl S, Neurath MF. 2017. Chemically induced mouse models of acute and chronic intestinal inflammation. Nat Protoc 12:1295–1309.

41. Eichele DD, Kharbanda KK. 2017. Dextran sodium sulfate colitis murine model: An indispensable tool for advancing our understanding of inflammatory bowel diseases pathogenesis. World J Gastroenterol 23:6016–6029.

42. Gong W, Yu J, Zheng T, Liu P, Zhao F, Liu J, Hong Z, Ren H, Gu G, Wang G, Wu X, Zhao Y, Ren J. 2021. CCL4-mediated targeting of spleen tyrosine kinase (Syk) inhibitor using nanoparticles alleviates inflammatory bowel disease. Clin Transl Med 11:e339.

43. Sikorska-Zimny K, Beneduce L. 2021. The Metabolism of Glucosinolates by Gut Microbiota. Nutrients 13:2750.

44. Luang-In V, Narbad A, Cebeci F, Bennett M, Rossiter JT. 2015. Identification of Proteins Possibly Involved in Glucosinolate Metabolism in *L. agilis* R16 and *E. coli* VL8. Protein J 34:135–146.

45. Chénard T, Malick M, Dubé J, Massé E. 2020. The influence of blood on the human gut microbiome. BMC Microbiol 20:44.

46. Cafiero C, Re A, Pisconti S, Trombetti M, Perri M, Colosimo M, D’Amato G, Gallelli L, Cannataro R, Molinario C, Fazio A, Caroleo MC, Cione E. 2020. Dysbiosis in intestinal microbiome linked to fecal blood determined by direct hybridization. 3 Biotech 10:358.

47. Chassaing B, Srinivasan G, Delgado MA, Young AN, Gewirtz AT, Vijay-Kumar M. 2012. Fecal Lipocalin 2, a Sensitive and Broadly Dynamic Non-Invasive Biomarker for Intestinal Inflammation. PLoS One 7:e44328.

48. Bábíčková J, Tóthová Ľ, Lengyelová E, Bartoňová A, Hodosy J, Gardlík R, Celec P. 2015. Sex Differences in Experimentally Induced Colitis in Mice: a Role for Estrogens. Inflammation 38:1996–2006.

49. Goodman WA, Havran HL, Quereshy HA, Kuang S, De Salvo C, Pizarro TT. 2018. Estrogen Receptor α Loss-of-Function Protects Female Mice From DSS-Induced Experimental Colitis. Cell Mol Gastroenterol Hepatol 5:630–633.e1.

50. Letourneau J, Holmes ZC, Dallow EP, Durand HK, Jiang S, Carrion VM, Gupta SK, Mincey AC, Muehlbauer MJ, Bain JR, David LA. 2022. Ecological memory of prior nutrient exposure in the human gut microbiome. ISME J 16:2479–2490.

51. Angelino D, Dosz EB, Sun J, Hoeflinger JL, Van Tassell ML, Chen P, Harnly JM, Miller MJ, Jeffery EH. 2015. Myrosinase-dependent and -independent formation and control of isothiocyanate products of glucosinolate hydrolysis. Front Plant Sci 6:831.

52. Stackebrandt E, Kramer I, Swiderski J, Hippe H. 1999. Phylogenetic basis for a taxonomic dissection of the genus *Clostridium*. FEMS Immunol Med Microbiol 24:253–258.

53. Uzal FA, McClane BA, Cheung JK, Theoret J, Garcia JP, Moore RJ, Rood JI. 2015. Animal models to study the pathogenesis of human and animal *Clostridium perfringens* infections. Vet Microbiol 179:23–33.

54. Chen H, Ma X, Liu Y, Ma L, Chen Z, Lin X, Si L, Ma X, Chen X. 2019. Gut Microbiota Interventions With *Clostridium butyricum* and Norfloxacin Modulate Immune Response in Experimental Autoimmune Encephalomyelitis Mice. Front Immunol 10:1662.

55. Shahinozzaman M, Raychaudhuri S, Fan S, Obanda DN. 2021. Kale Attenuates Inflammation and Modulates Gut Microbial Composition and Function in C57BL/6J Mice with Diet-Induced Obesity. Microorganisms 9:238.

56. Ju T, Kong JY, Stothard P, Willing BP. 2019. Defining the role of *Parasutterella*, a previously uncharacterized member of the core gut microbiota. ISME J 13:1520–1534.

57. Hubbard TD, Murray IA, Nichols RG, Cassel K, Podolsky M, Kuzu G, Tian Y, Smith P, Kennett MJ, Patterson AD, Perdew GH. 2017. Dietary Broccoli Impacts Microbial Community Structure and Attenuates Chemically Induced Colitis in Mice in an Ah receptor dependent manner. J Funct Foods 37:685–698.

58. Antonson AM, Evans MV, Galley JD, Chen HJ, Rajasekera TA, Lammers SM, Hale VL, Bailey MT, Gur TL. 2020. Unique maternal immune and functional microbial profiles during prenatal stress. Sci Rep 10:20288.

59. Huarte-Mendicoa JC, Astiasarán I, Bello J. 1997. Nitrate and nitrite levels in fresh and frozen broccoli. Effect of freezing and cooking. Food Chem 58:39–42.

60. Araki Y, Mukaisho K, Sugihara H, Fujiyama Y, Hattori T. 2010. *Proteus mirabilis* sp. intestinal microflora grow in a dextran sulfate sodium-rich environment. Int J Mol Med 25:203–208.

61. Chang C-S, Liao Y-C, Huang C-T, Lin C-M, Cheung CHY, Ruan J-W, Yu W-H, Tsai Y-T, Lin I-J, Huang C-H, Liou J-S, Chou Y-H, Chien H-J, Chuang H-L, Juan H-F, Huang H-C, Chan H-L, Liao Y-C, Tang S-C, Su Y-W, Tan T-H, Bäumler AJ, Kao C-Y. 2021. Identification of a gut microbiota member that ameliorates DSS-induced colitis in intestinal barrier enhanced Dusp6-deficient mice. Cell Rep 37:110016.

62. Khan I, Bai Y, Ullah N, Liu G, Rajoka MSR, Zhang C. 2022. Differential Susceptibility of the Gut Microbiota to DSS Treatment Interferes in the Conserved Microbiome Association in Mouse Models of Colitis and Is Related to the Initial Gut Microbiota Difference. Advanced Gut & Microbiome Research 2022:7813278.

63. Gibbons SM. 2020. Keystone taxa indispensable for microbiome recovery. Nature Microbiology 5:1067–1068.

64. Duvallet C, Gibbons SM, Gurry T, Irizarry RA, Alm EJ. 2017. Meta-analysis of gut microbiome studies identifies disease-specific and shared responses. Nat Commun 8:1784.

65. Walters WA, Xu Z, Knight R. 2014. Meta-analyses of human gut microbes associated with obesity and IBD. FEBS Lett 588:4223–4233.

66. Han YW. 2015. *Fusobacterium nucleatum*: a commensal-turned pathogen. Curr Opin Microbiol 23:141–147.

67. Tropini C, Earle KA, Huang KC, Sonnenburg JL. 2017. The Gut Microbiome: Connecting Spatial Organization to Function. Cell Host Microbe 21:433–442.

68. Gu M, Samuelson DR, de la Rua NM, Charles TP, Taylor CM, Luo M, Siggins RW, Shellito JE, Welsh DA. 2022. Host innate and adaptive immunity shapes the gut microbiota biogeography. Microbiol Immunol 66:330–341.

69. Suzuki TA, Nachman MW. 2016. Spatial Heterogeneity of Gut Microbial Composition along the Gastrointestinal Tract in Natural Populations of House Mice. PLoS One 11:e0163720.

70. Wu S, Zhang B, Liu Y, Suo X, Li H. 2018. Influence of surface topography on bacterial adhesion: A review (Review). Biointerphases 13:060801.

71. Stenman LK, Holma R, Forsgård R, Gylling H, Korpela R. 2013. Higher fecal bile acid hydrophobicity is associated with exacerbation of dextran sodium sulfate colitis in mice. J Nutr 143:1691–1697.

72. Fahey JW, Smilovitz Burak J, Evans D. 2022. Sprout microbial safety: A reappraisal after a quarter-century. Food Frontiers 1–7.

73. Rao M, Afshin A, Singh G, Mozaffarian D. 2013. Do healthier foods and diet patterns cost more than less healthy options? A systematic review and meta-analysis. BMJ Open 3:e004277.

74. Economic Research Service. Food Access Research Atlas. US Department of Agriculture. https://www.ers.usda.gov/data/fooddesert. Retrieved 22 April 2022.

75. Wedick NM, Ma Y, Olendzki BC, Procter-Gray E, Cheng J, Kane KJ, Ockene IS, Pagoto SL, Land TG, Li W. 2015. Access to healthy food stores modifies effect of a dietary intervention. Am J Prev Med 48:309–317.

76. Baker EA, Schootman M, Barnidge E, Kelly C. 2006. The role of race and poverty in access to foods that enable individuals to adhere to dietary guidelines. Prev Chronic Dis 3:A76.

77. Inotiv. TD.97184: AIN93G Diet. Inotiv. https://insights.envigo.com/hubfs/resources/data-sheets/97184.pdf. Retrieved 28 April 2023.

78. Mähler M, Bristol IJ, Leiter EH, Workman AE, Birkenmeier EH, Elson CO, Sundberg JP. 1998. Differential susceptibility of inbred mouse strains to dextran sulfate sodium-induced colitis. Am J Physiol 274:G544–51.

79. Okayasu I, Hatakeyama S, Yamada M, Ohkusa T, Inagaki Y, Nakaya R. 1990. A novel method in the induction of reliable experimental acute and chronic ulcerative colitis in mice. Gastroenterology 98:694–702.

80. Poritz LS, Garver KI, Green C, Fitzpatrick L, Ruggiero F, Koltun WA. 2007. Loss of the tight junction protein ZO-1 in dextran sulfate sodium induced colitis. J Surg Res 140:12–19.

81. Chassaing B, Aitken JD, Malleshappa M, Vijay-Kumar M. 2014. Dextran sulfate sodium (DSS)-induced colitis in mice. Curr Protoc Immunol 104:15.25.1–15.25.14.

82. Araki A, Kanai T, Ishikura T, Makita S, Uraushihara K, Iiyama R, Totsuka T, Takeda K, Akira S, Watanabe M. 2005. MyD88-deficient mice develop severe intestinal inflammation in dextran sodium sulfate colitis. J Gastroenterol 40:16–23.

83. Dieleman LA, Ridwan BU, Tennyson GS, Beagley KW, Bucy RP, Elson CO. 1994. Dextran sulfate sodium-induced colitis occurs in severe combined immunodeficient mice. Gastroenterology 107:1643–1652.

84. Taghipour N, Molaei M, Mosaffa N, Rostami-Nejad M, Asadzadeh Aghdaei H, Anissian A, Azimzadeh P, Zali MR. 2016. An experimental model of colitis induced by dextran sulfate sodium from acute progresses to chronicity in C57BL/6: correlation between conditions of mice and the environment. Gastroenterol Hepatol Bed Bench 9:45–52.

85. Yu W, Li Q, Shao C, Zhang Y, Kang C, Zheng Y, Liu X, Liu X, Yan J. 2022. The Cao-Xiang-Wei-Kang formula attenuates the progression of experimental colitis by restoring the homeostasis of the microbiome and suppressing inflammation. Front Pharmacol 13:946065.

86. Prakash T, Janadri S. 2022. Anti-inflammatory effect of wedelolactone on DSS induced colitis in rats: IL-6/STAT3 signaling pathway. J Ayurveda Integr Med 100544.

87. Gobert AP, Al-Greene NT, Singh K, Coburn LA, Sierra JC, Verriere TG, Luis PB, Schneider C, Asim M, Allaman MM, Barry DP, Cleveland JL, Destefano Shields CE, Casero RA Jr, Washington MK, Piazuelo MB, Wilson KT. 2018. Distinct Immunomodulatory Effects of Spermine Oxidase in Colitis Induced by Epithelial Injury or Infection. Front Immunol 9:1242.

88. Nocker A, Sossa-Fernandez P, Burr MD, Camper AK. 2007. Use of propidium monoazide for live/dead distinction in microbial ecology. Appl Environ Microbiol 73:5111–5117.

89. The Earth Microbiome Project. 2022. The Earth Microbiome Project. The Earth Microbiome Project. https://earthmicrobiome.org/. Retrieved 22 April 2022.

90. Callahan BJ, McMurdie PJ, Rosen MJ, Han AW, Johnson AJA, Holmes SP. 2016. DADA2: High-resolution sample inference from Illumina amplicon data. Nat Methods 13:581–583.

91. RCoreTeam. 2022. R: a language and environment for statistical computing (4.2.2). R Foundation for Statistical Computing, Vienna, Austria. https://www.r-project.org/.

92. Pruesse E, Quast C, Knittel K, Fuchs BM, Ludwig W, Peplies J, Glöckner FO. 2007. SILVA: a comprehensive online resource for quality checked and aligned ribosomal RNA sequence data compatible with ARB. Nucleic Acids Res 35:7188–7196.

93. Wickham H, Francois R, Henry L, Müller K. 2015. dplyr: A Grammar of Data Manipulation. R package version 0.4. 3. R Found Stat Comput, Vienna https://CRANRprojectorg/package=dplyr.

94. Ishaq SL. 2017. Phyloseq_dealing_with_neg_controls_ Ishaq_example.R. Github. https://github.com/SueIshaq/Examples-DADA2-Phyloseq. Retrieved 5 March 2019.

95. McMurdie PJ, Holmes S. 2013. phyloseq: An R Package for Reproducible Interactive Analysis and Graphics of Microbiome Census Data. PLoS One 8:e61217.

96. Bates D, Mächler M, Zurich E, Bolker BM, Walker SC. 2015. Fitting Linear Mixed-Effects Models Using lme4. Journal of Statistical Software, Articles 67:1–48.

97. Pedersen EJ, Miller DL, Simpson GL, Ross N. 2019. Hierarchical generalized additive models: an introduction with mgcv. PeerJ 7:e6876.

98. Oksanen J, Guillaume Blanchet F, Friendly M, Kindt R, Legendre P, McGlinn D, Minchin PR, O’Hara RB, Simpson GL, Solymos P, Stevens MHH, Szoecs E, Wagner H. 2020. Vegan: Community Ecology Package (2.5-7). https://cran.r-project.org/web/packages/vegan/vignettes/diversity-vegan.pdf.

99. Archer E. 2022. rfpermute: Estimate Permutation p-Values for Random Forest Importance Metrics (2.5.1). https://github.com/EricArcher/rfPermute.

100. Wickham H. 2016. ggplot2: Elegant graphics for Data Analysis. Springer Publishing Company, New York. https://dl.acm.org/citation.cfm?id=1795559.

101. Kassambara A. 2022. ggpubr:“ggplot2” based publication ready plots (0.1.7). https://rpkgs.datanovia.com/ggpubr/.

102. Knights D, Kuczynski J, Charlson ES, Zaneveld J, Mozer MC, Collman RG, Bushman FD, Knight R, Kelley ST. 2011. Bayesian community-wide culture-independent microbial source tracking. Nat Methods 8:761–763.

103. Galey M. 2021. Mi-Seq Analysis. GitHub. https://mgaley-004.github.io/MiSeq-Analysis/Tutorials/SourceSink.html, https://github.com/mgaley-004/MiSeq-Analysis. Retrieved 7 December 2022.

