## Supplemental Figures and Table for "Steamed broccoli sprouts alleviate DSS-induced inflammation and retain gut microbial biogeography in mice"

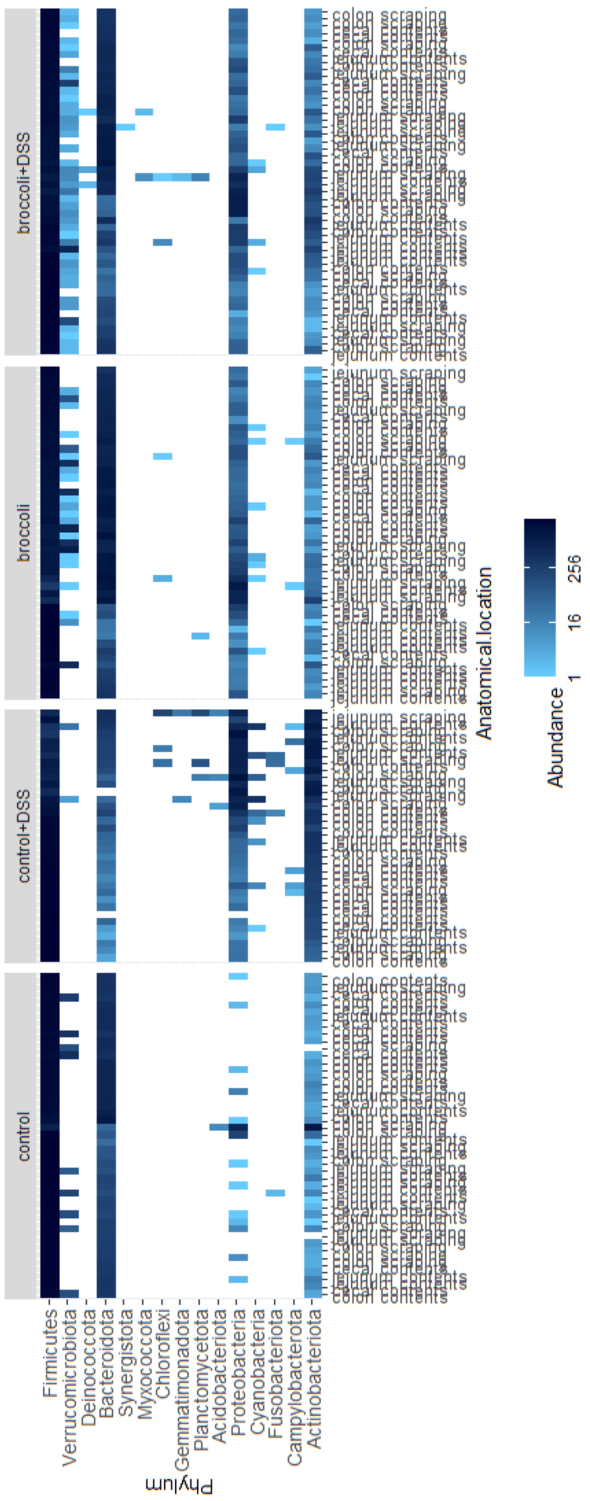


**Figure S1. Abundance of bacterial sequence reads identified at the phylum level, paneled by Treatment and labeled by anatomical location of the sample in the gastrointestinal tract of mice.** Four treatment groups were used in a 34-day chronic relapsing model of colitis: control diet, control diet with DSS added to drinking water, control diet adjusted with 10% by weight steamed broccoli sprouts, and 10% broccoli sprout diet with DSS added to drinking water (Figure 1). Bacterial communities were sampled from several locations in the gastrointestinal tract at the end of the study.

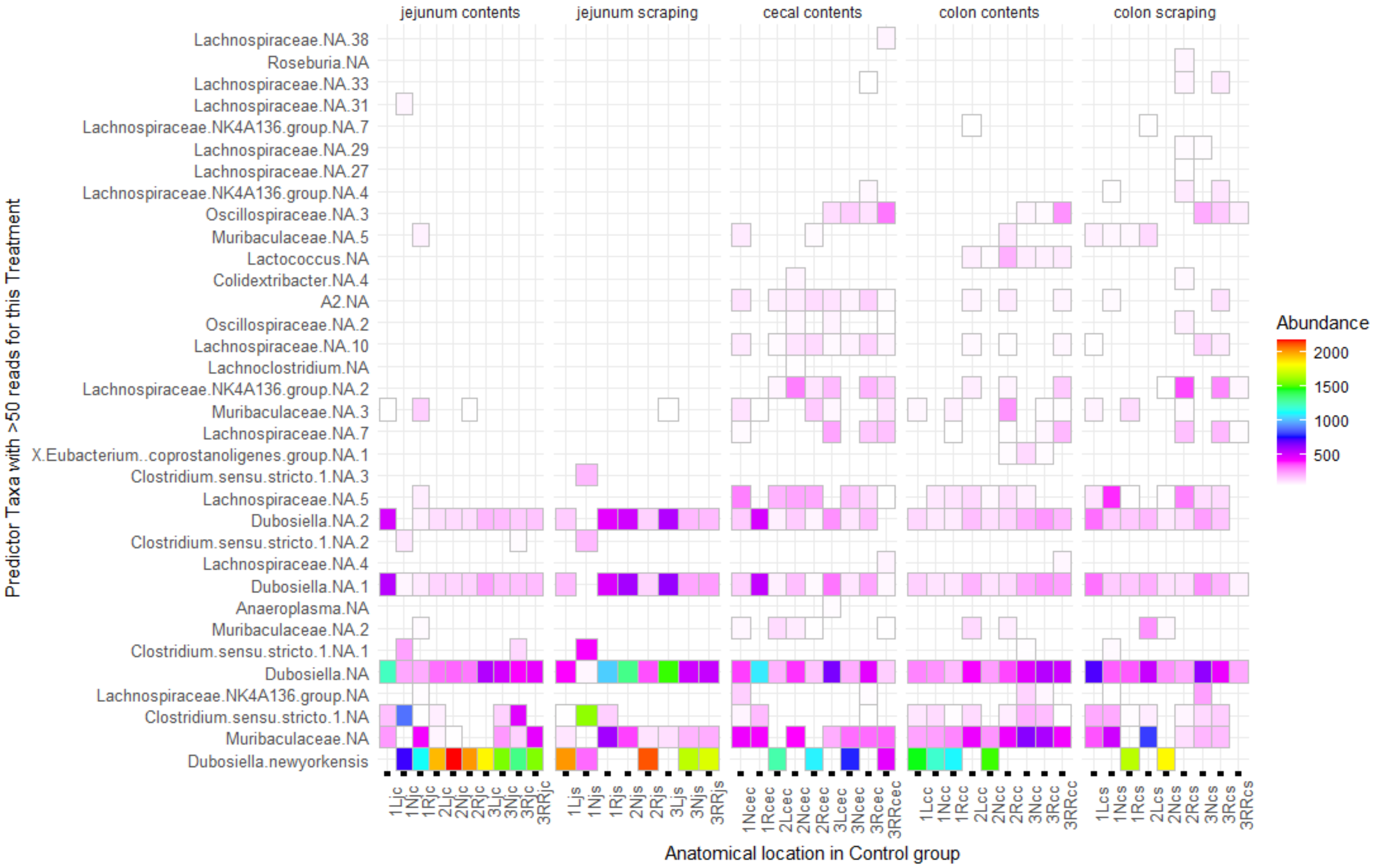


**Figure S2. Abundance of bacterial sequence variants identified as differential for the Control group**. Important features (SVs) were identified through permutational random forest analysis, and only the features important to this group (>50 reads) are listed out of 188 significant ($p < 0.05$) features across all treatments. Model accuracy was 98%. Bacterial sequence variants (SV) are identified as the lowest level of taxonomic identity possible, with "NA" indicating which could not be identified to species, and the number indicating which specific SV it was. Four treatment groups were used in a 34-day chronic relapsing model of colitis: control diet, control diet with DSS added to drinking water, control diet adjusted with 10% by weight steamed broccoli sprouts, and 10% broccoli sprout diet with DSS added to drinking water (Figure 1). Bacterial communities were sampled from several locations in the gastrointestinal tract at the end of the study.

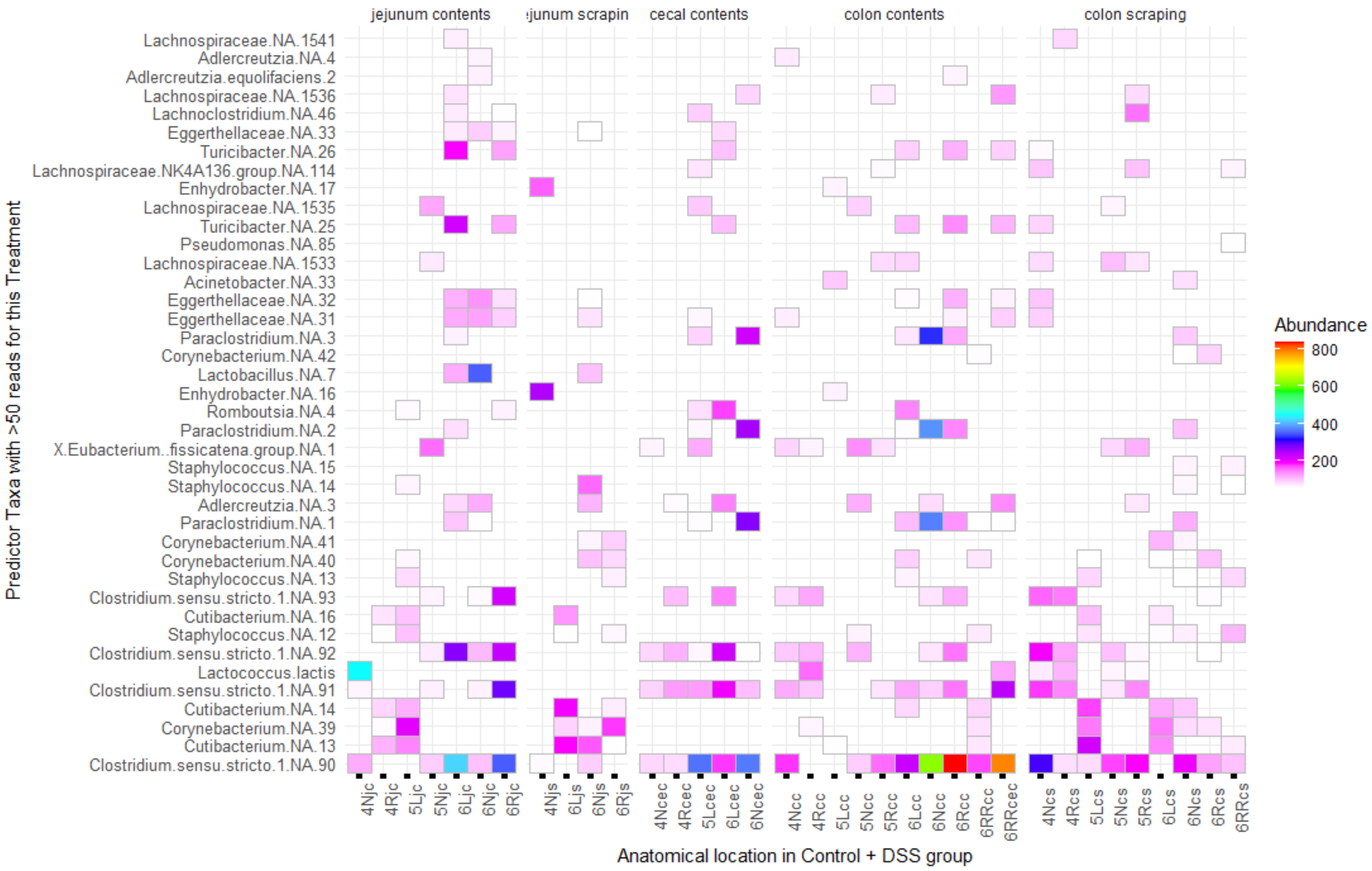


**Figure S3. Abundance of bacterial sequence variants identified as differential for the Control+DSS group**. Important features (SVs) were identified through permutational random forest analysis, and only the features important to this group (>50 reads) are listed out of 188 significant ($p < 0.05$) features across all treatments. Model accuracy was 98%. Bacterial sequence variants (SV) are identified as the lowest level of taxonomic identity possible, with "NA" indicating which could not be identified to species, and the number indicating which specific SV it was. Four treatment groups were used in a 34-day chronic relapsing model of colitis: control diet, control diet with DSS added to drinking water, control diet adjusted with 10% by weight steamed broccoli sprouts, and 10% broccoli sprout diet with DSS added to drinking water (Figure 1). Bacterial communities were sampled from several locations in the gastrointestinal tract at the end of the study.

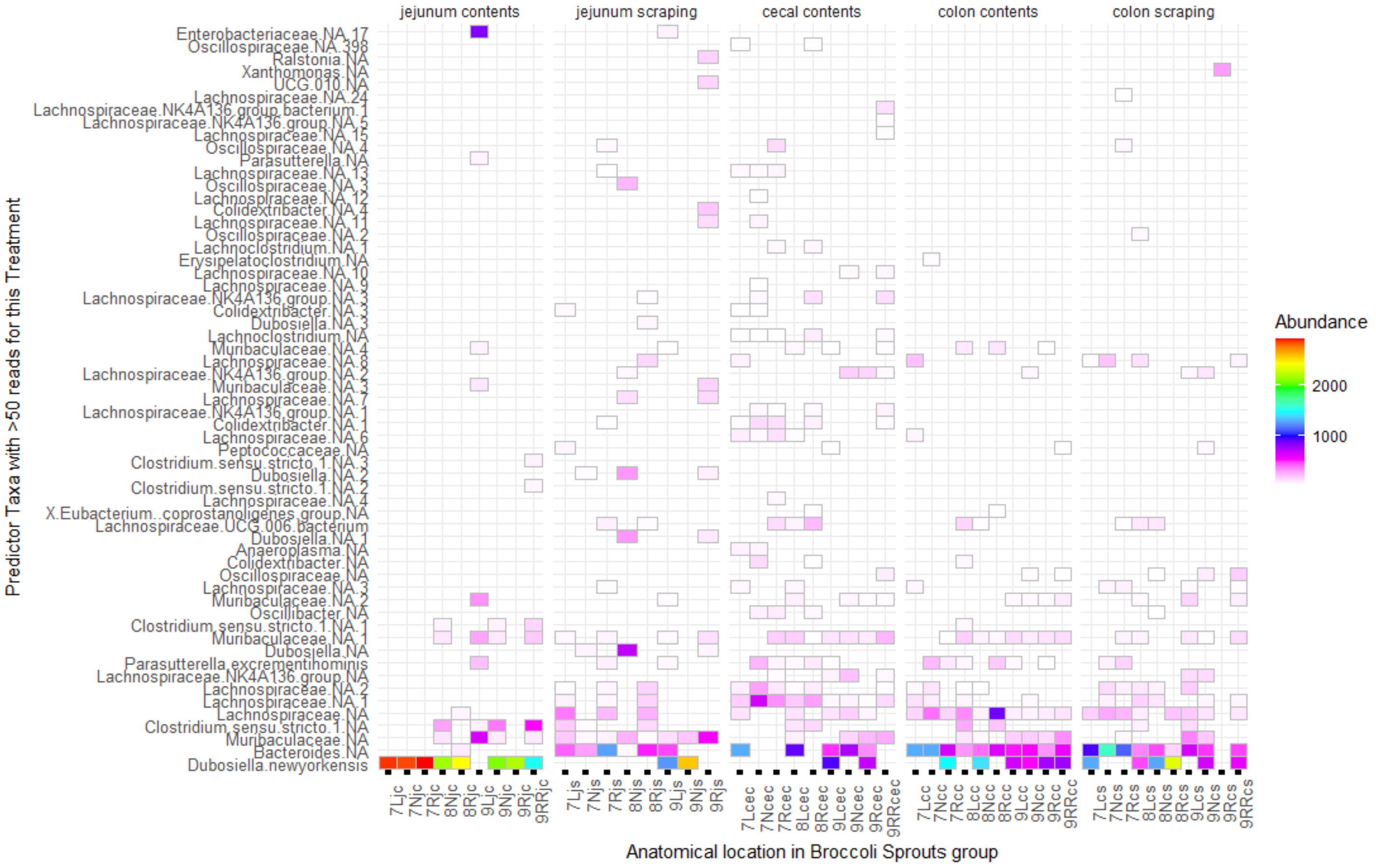


**Figure S4. Abundance of bacterial SVs identified as differential for the 10% steamed Broccoli Sprouts group**. Important features (SVs) were identified through permutational random forest analysis, and only the features important to this group (>50 reads) are listed out of 188 significant ($p < 0.05$) features across all treatments. Model accuracy was 98%. Bacterial sequence variants (SV) are identified as the lowest level of taxonomic identity possible, with “NA” indicating which could not be identified to species, and the number indicating which specific SV it was. Four treatment groups were used in a 40-day chronic relapsing model of colitis: control diet, control diet with DSS added to drinking water, control diet adjusted with 10% by weight steamed broccoli sprouts, and 10% broccoli sprout diet with DSS added to drinking water (Figure 1). Bacterial communities were sampled from several locations in the gastrointestinal tract at the end of the study.

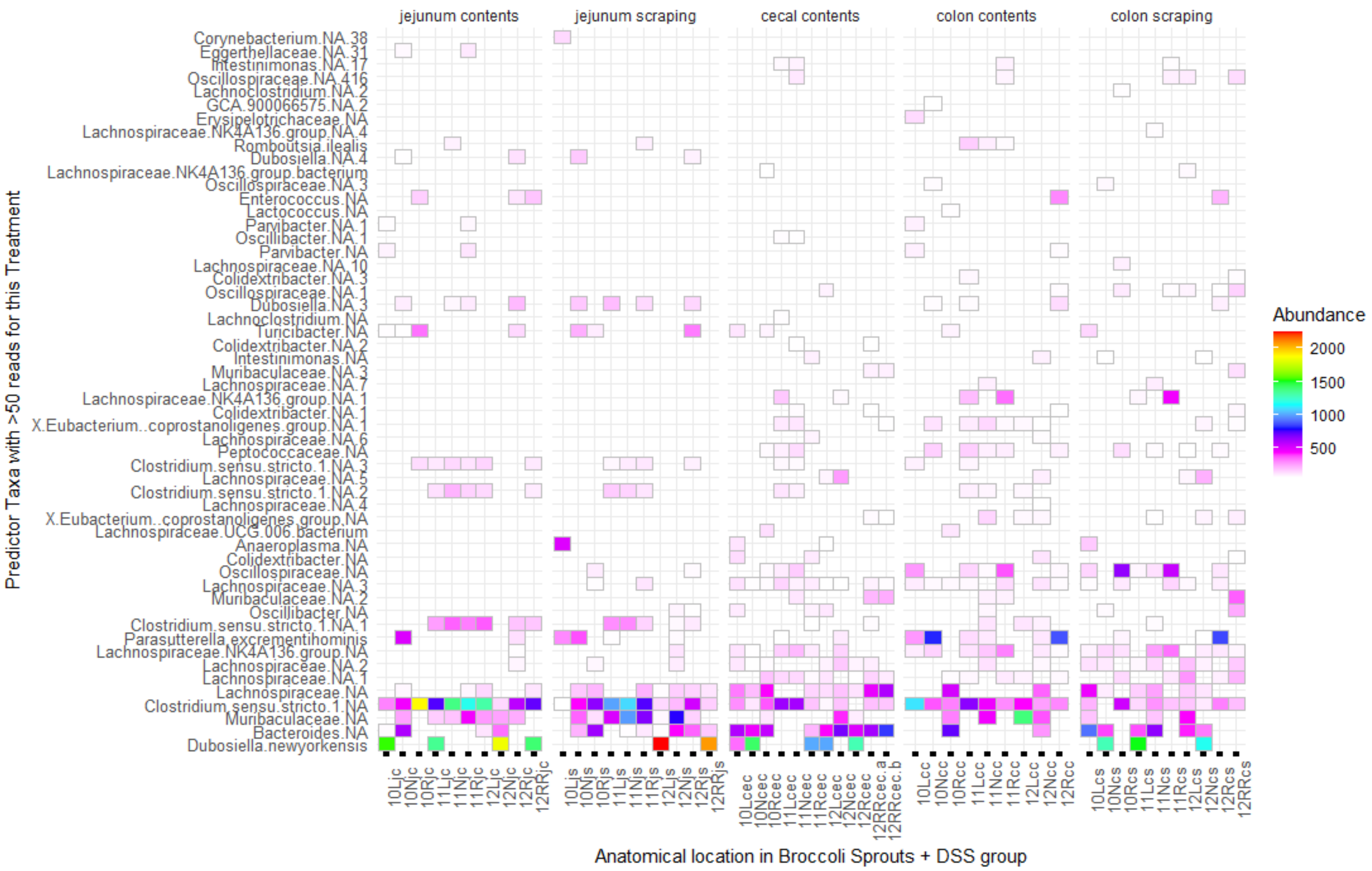


**Figure S5. Abundance of bacterial SVs identified as differential for the Broccoli+DSS group**. Important features (SVs) were identified through permutational random forest analysis, and only the features important to this group (>50 reads) are listed out of 188 significant ($p < 0.05$) features across all treatments. Model accuracy was 98%. Bacterial sequence variants (SV) are identified as the lowest level of taxonomic identity possible, with "NA" indicating which could not be identified to species, and the number indicating which specific SV it was. Four treatment groups were used in a 34-day chronic relapsing model of colitis: control diet, control diet with DSS added to drinking water, control diet adjusted with 10% by weight steamed broccoli sprouts, and 10% broccoli sprout diet with DSS added to drinking water (Figure 1). Bacterial communities were sampled from several locations in the gastrointestinal tract at the end of the study.

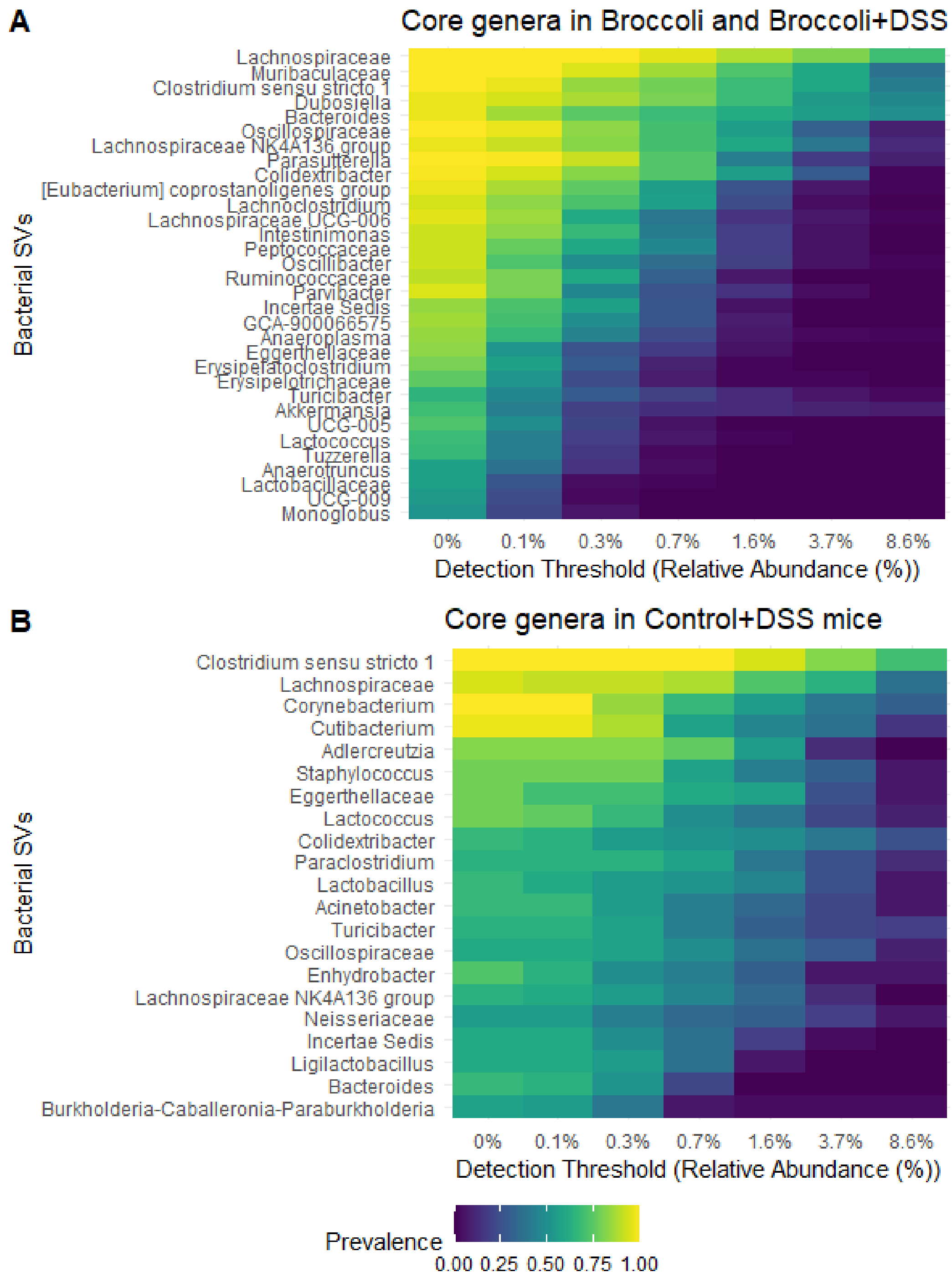


**Figure S6. Core bacterial sequence variants (SVs) shared by 70% of gut samples from A) mice consuming a 10% steamed broccoli sprout diet with or without DSS added to drinking water, or B) mice consuming a control diet with DSS in drinking water.** There were no bacterial SVs shared across 70% of Control+DSS and Broccoli+DSS samples (data not shown). Four treatment groups were used in a 34-day chronic relapsing model of colitis: control diet, control diet with DSS added to drinking water, control diet adjusted with 10% by weight steamed broccoli sprouts, and 10% broccoli sprout diet with DSS added to drinking water (Figure 1). Bacterial communities were sampled from several locations in the gastrointestinal tract at the end of the study.

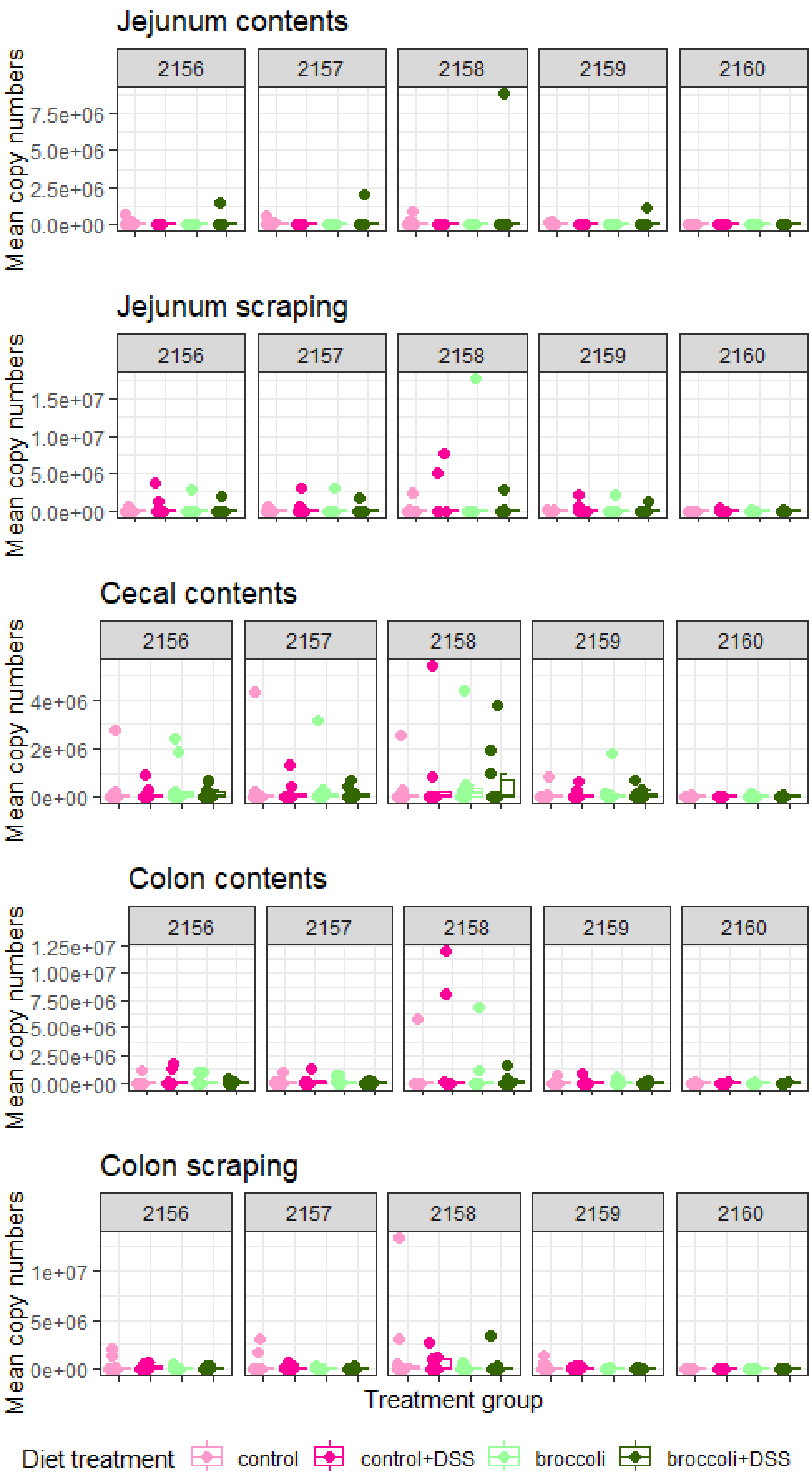


**Figure S7: qPCR results for the Bacterial operon BT2160-BT2156 for *Bacteroides thetaiotaomicron* in mice under a DSS-induced model of chronic, relapsing colitis with or without 10% steamed broccoli sprouts in diet.**

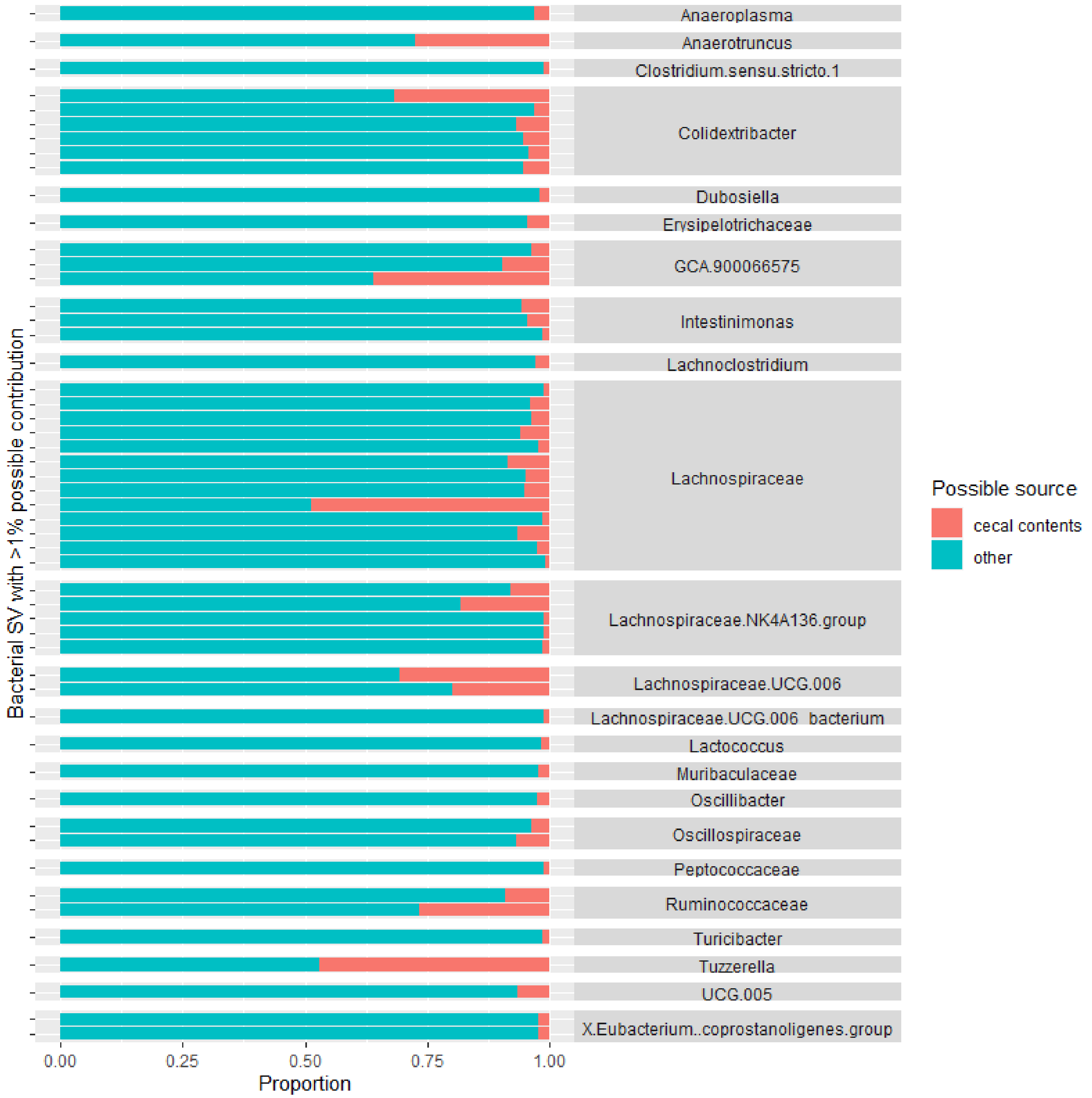


**Figure S8. Bacterial sequence variants which are hypothesized to be sourced from the cecum and creating population sinks in other locations in the intestines of mice.** Potential source SVs were identified with the SourceTracker algorithm modified for the R platform. A total of 142 SVs were identified, and 95 had a proportion >1% and are visualized here. Four treatment groups were used in a 34-day chronic relapsing model of colitis: control diet, control diet with DSS added to drinking water, control diet adjusted with 10% by weight steamed broccoli sprouts, and 10% broccoli sprout diet with DSS added to drinking water (Figure 1). Bacterial communities were sampled from several locations in the gastrointestinal tract at the end of the study.

**Supplemental Tables**

**Table S1 Targets for quantitative polymerase chain reaction (qPCR) of glucosinolate metabolizing enzymes in *Bacteroidetes thetaiotaomicron*.**

| Set | Forward Primer | Reverse Primer | Amplicon size, gene | Reference |
|---|---|---|---|---|
| BT2160 | BT2160_fwd:<br>CAGAATGCCACA<br>CGCGAAAC | BT2160_rev:<br>TGAATGCCGGA<br>GGAAACGAC | 127 bp, SusR4,<br>Transcriptional regulator protein on inner membrane. Controls BT2159-2156. | (Liou et al. 2020) |
| Gene block | TGCTGATATTCAGAATGCCACACGCGAAACAGCCTCCCTGCAGGCACTTGCACTG<br>ATACAATATGAGGAAAATAATTTAGCGGATGCTTTCAAATTTACTCAGTCGGCTAT<br>TGATGATGTCGTTTCCTCCGGCATTCATTTCCGGGCA | | | |
| Protocol | 1) 1 cycle at 50°C for 2 min; 2) 1 cycle at 95°C for 1 min; 3) 40 cycles at 95°C for 15 s, 60°C for 30 s and 72°C for 30 s, followed by a plate read, | | | |
| **Set** | **Forward Primer** | **Reverse Primer** | **Amplicon size, gene** | **Reference** |
| BT2159 | BT2159_ fwd<br>TGCGATACAGAT<br>CCTACCACGC | BT2159_rev<br>CAATGTGAAGA<br>GCCCGACAACC | 123 bp, Nicotinamide-dependent oxidoreductase. Cytoplasmic. | (Liou et al. 2020) |
| Gene block | AGATTGCCTATATTTGCGATACAGATCCTACCACGCGCGAACTTGCTAAAAAAAT<br>ATCTCCTGAATCTCTTGTAGTAGAGAATGATCAAAAAATCTTTGAAGACGAGAGT<br>GTACAGGTTGTCGGGCTCTTCACATTGGCAGACTCAAG | | | |
| Protocol | 1) 1 cycle at 50°C for 2 min; 2) 1 cycle at 95°C for 1 min; 3) 40 cycles at 95°C for 15 s, 60°C for 30 s and 72°C for 30 s, followed by a plate read, | | | |
| **Set** | **Forward Primer** | **Reverse Primer** | **Amplicon size, gene** | **Reference** |
| BT2158 | BT2158_fwd<br>CGAAACAATTTG<br>CAGCCGAAC | BT2158_rev<br>GGCAACTTCCA<br>TCCTTCACG | 58 bp, Nicotinamide-dependent oxidoreductase. Periplasmic. | (Liou et al. 2020) |
| Gene block | TTCATGATGGTCACCCGTCATTCAATAAGACCTGGACAGATCCGATCAA<br>CGCGAAACAATTTGCAGCCGAACTGGTGAAACATAATTATCGTGAAGGATGGAAG<br>TTGCCTGATATGCCACGATAA | | | |
| Protocol | 1) 1 cycle at 50°C for 2 min; 2) 1 cycle at 95°C for 1 min; 3) 40 cycles at 95°C for 15 s, 60°C for 30 s and 72°C for 20 s, followed by a plate read, | | | |
| **Set** | **Forward Primer** | **Reverse Primer** | **Amplicon size, gene** | **Reference** |
| BT2157 | BT2157_fwd<br>TGCAAGCCAGCA<br>AATTCAGC | BT2157_rev<br>CAGTCCAGAAC<br>TTTCACGCG | 151 bp, Glycosyl hydrolase. Outer membrane lipoprotein. | (Liou et al. 2020) |
| Gene block | ACCTTTTGCAAGCCAGCAAATTCAGCCAGGACAAATGGCCGTTGGCTTTCGAACT<br>GCTGAATAATTGCGGTGGCGAAAACCACGAAGGATTTATCGGAATGCAGGATCAC<br>GGTGATGACGTTTGGTTCCGCAATATCCGCGTGAAAGTTCTGGACTGA | | | |
| Protocol | 1) 1 cycle at 50°C for 2 min; 2) 1 cycle at 95°C for 1 min; 3) 40 cycles at 95°C for 15 s, 60°C for 30 s and 72°C for 30 s, followed by a plate read, | | | |
| **Set** | **Forward Primer** | **Reverse Primer** | **Amplicon size, gene** | **Reference** |
| BT2156 | BT2156_fwd<br>CTGCCGGGCTGA<br>AGGTTTTATC | BT2156_rev<br>TCAGCAATACAC<br>TGGTCCCACC | 112 bp, Sugar phosphate isomerase. | (Liou et al. 2020) |
| Gene block | TGTTGAATCTGCCGGGCTGAAGGTTTTATCCTCACATTGCACAAGAGGATTGTCGA<br>AAGAAGAATTAGCTTCCGGTGATTTTTCAAGTTCACTTCAATGGTGGGACCAGTGT<br>ATTGCTGATCATA | | | |
| Protocol | 1) 1 cycle at 50°C for 2 min; 2) 1 cycle at 95°C for 1 min; 3) 40 cycles at 95°C for 15 s, 60°C for 25 s and 72°C for 25 s, followed by a plate read, | | | |
